# Engineering de novo binder CAR-T cell therapies with generative AI

**DOI:** 10.1101/2024.11.25.625151

**Authors:** Markus Mergen, Daniela Abele, Naile Koleci, Alba Schmahl Fernandez, Maya Sugden, Noah Holzleitner, Andreas Carr, Leonie Rieger, Valentina Leone, Maximilian Reichert, Karl-Ludwig Laugwitz, Florian Bassermann, Dirk H. Busch, Julian Grünewald, Andrea Schmidts

**Affiliations:** Department of Medicine III, Hematology and Oncology, TUM University Hospital, Technical University of Munich, TUM School of Medicine and Health, Munich, Germany; Center for Translational Cancer Research (TranslaTUM), Technical University of Munich, Munich, Germany; German Cancer Consortium (DKTK), partner-site Munich, a partnership between DKFZ and Klinikum rechts der Isar, Munich, Germany; Department of Medicine I, Cardiology, Angiology and Pneumology, TUM University Hospital, Technical University of Munich, TUM School of Medicine and Health, Munich, Germany; TUM Center for Organoid Systems and Tissue Engineering (COS), Garching, Germany; Institute for Medical Microbiology, Immunology and Hygiene, Technical University of Munich, TUM School of Medicine and Health, Munich, Germany; German Center for Infection Research (DZIF), Partner Site Munich, Munich, Germany; Translational Pancreatic Cancer Research Center, TUM School of Medicine and Health, Department of Medicine II, Technical University of Munich, Munich, Germany; Department of Medicine II, Endocrinology, Diabetology, Infectious Diseases and Gastroenterology, TUM University Hospital, Technical University of Munich, TUM School of Medicine and Health, Munich, Germany; Center for Protein Assemblies (CPA), Technical University of Munich, Garching, Germany; Bavarian Cancer Research Center (BZKF), Munich, Germany; German Center of Cardiovascular Research (DZHK), Munich Heart Alliance, Munich, Germany

## Abstract

Chimeric antigen receptor T cell (CAR-T) therapies have revolutionized cancer treatment, with six CAR-T products currently in clinical use^1–4^. Despite their success, high resistance rates due to antigen escape remain a major challenge^5,6^. In silico design of de novo binders (DNBs) has the potential to accelerate the development of new binding domains for CAR-T, possibly enabling personalized therapies for cancer resistance^7,8^. Here, we show that DNBs can be used for CAR-T, targeting clinically relevant cancer antigens. Using a DNB against the epidermal growth factor receptor (EGFR), we demonstrate comparable cytotoxicity, cytokine secretion, long-term proliferation, and lysis of primary patient-derived cancer organoids with single-chain variable fragment (scFv)-based and DNB-based CAR-T cells. Moreover, we use generative artificial intelligence (AI) guided binder design with RFdiffusion^9^ to target the B cell maturation antigen (BCMA), a key antigen in multiple myeloma treatment^10–17^. We confirmed the activity of our AI-designed BCMA CAR-T in short- and long-term effector readouts, including a xenograft mouse model of multiple myeloma. Notably, our AI-guided CAR-T approach also successfully targets a mutated BCMA protein variant resistant to the clinically used bispecific antibody teclistamab. In sum, we demonstrate a proof-of-concept for engineering new, bespoke cellular immunotherapies targeting cancer resistance with the help of generative AI. This approach may further accelerate the development of new CAR-T therapies addressing cancer resistance.

## Introduction

Alternative types of antigen-binding domains have successfully been used for CAR designs, and several of them are currently being tested in early-phase clinical trials (NCT04661384, NCT02830724, NCT05020444)^18^. Most commonly, these binders either leverage naturally occurring receptor-ligand pairs or mimic antibody structures^19–21^. Recently, computational models were developed that enable the generation of de novo binders when provided with the target protein structure alone^7^. These were made possible by deep learning approaches that have greatly advanced both sequence-to-structure (Alphafold and Rosettafold) and structure-to-sequence predictions^22–24^. RoseTTAFold diffusion (RFdiffusion), a generative diffusion model for protein design, uses a combination of both approaches. Moreover, RFdiffusion is a universally accessible and easy-to-use method that holds promise for democratizing structure-based binder design^9^. The use of such generative AI-guided mini-binders could greatly accelerate the engineering of new CAR-T therapies, especially to target resistant tumors. While recent studies have begun to explore the potential of de novo binders within CAR architectures^25^, comprehensive evaluations of their functional performance across diverse target antigens, and clinical settings remain largely unaddressed. In this work, we use previously described de novo binders (DNB) to engineer and characterize both T cell engager (TCE) and CAR-T therapies side-by-side with clinically used single-chain variable fragment (scFv) binding domains. More importantly, we subsequently use RFdiffusion to generate a de novo binder targeting the B-cell maturation antigen (BCMA) and demonstrate the potential of AI-guided CAR-T therapies.

## Results

To test computationally generated DNBs in cellular immunotherapies, we initially investigated the activity of a previously established binder in a TCE (**Fig. 1a**). This DNB is a 62 amino acid (AA) triple-helix bundle targeting the N-terminal domain of EGFR, previously tested for in vitro binding only^7^. We tested the DNB in a T cell engaging (TCE) molecule, where it was fused to a CD3-binding scFv (DNB-TCE). This construct was tested side-by-side with an anti-EGFR TCE derived from the monoclonal antibody cetuximab (scFv-TCE) (**Extended Data Fig. 1a**). Both TCEs were produced in HEK293T cells and added to co-cultures of primary T cells and BxPC3 target cells that endogenously express EGFR. Subsequently, we measured dose-dependent binding of TCEs to tumor cells in flow cytometry as well as TCE-mediated tumor cell lysis using a luciferase-based readout (**Fig. 1b**, **Extended Data Fig. 1b,** and **Supplementary Table 1**). Lysis induced by both DNB- and scFv-based TCEs was similar across all tested effector-to-target (E:T) ratios (**Fig. 1b**).

**Fig. 1.**
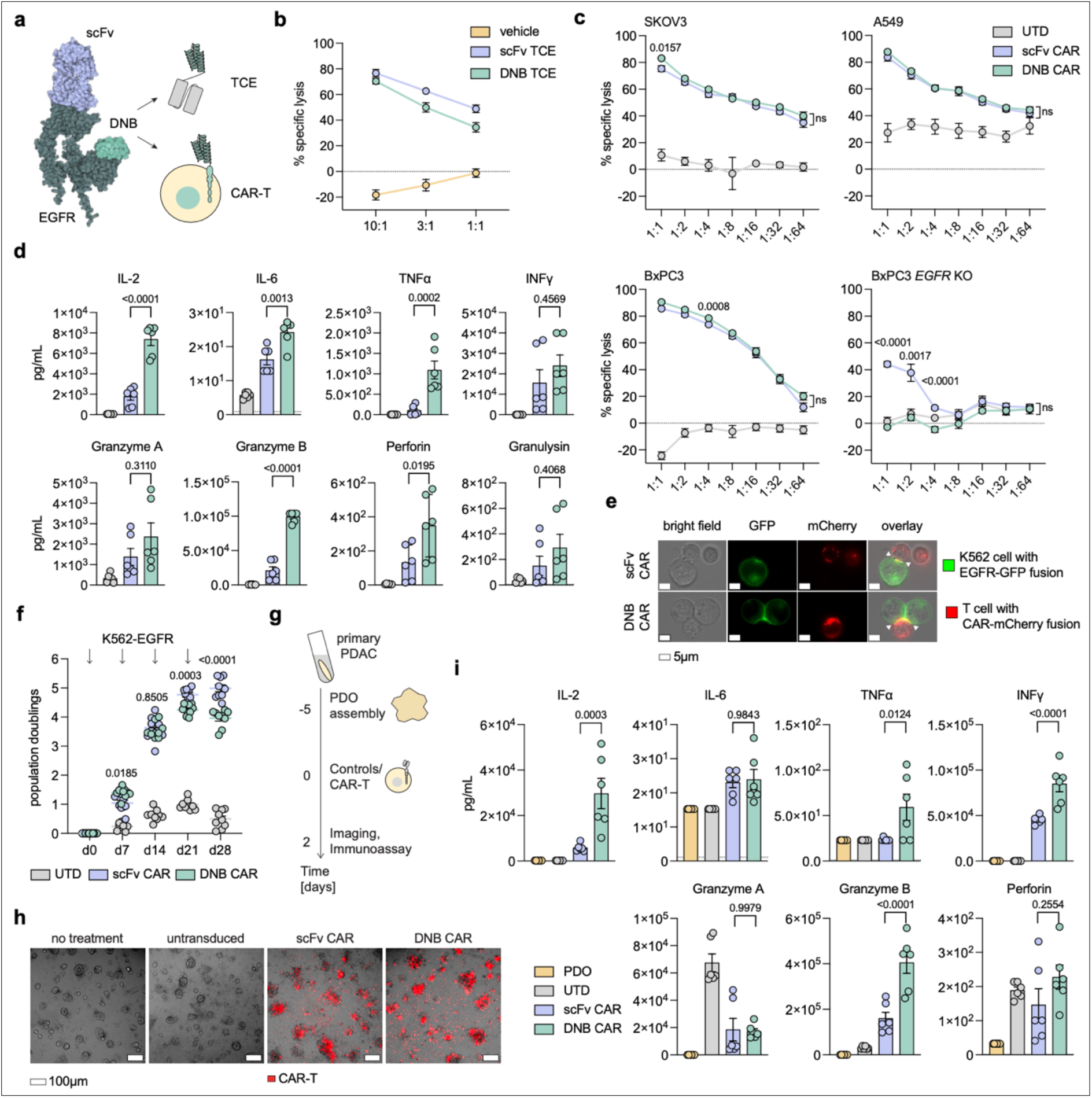
Engineering and characterization of cellular immunotherapies using de novo binders. **a**, Schematic comparing a cetuximab-derived scFv (violet) to a DNB (green)^7^, both binding the extracellular domain of EGFR (gray) at different sites (left panel). Hypothesized use of DNBs for TCE and CAR-T therapies (right panel; not drawn to scale). **b,** Cytotoxicity mediated by DNB and scFv TCE against tumor target BxPC3 cells in the presence of primary T cells at indicated effector:target (E:T) ratios over 18h of co-culture. **c**, Cytotoxicity assays of untransduced T cells (UTD), DNB, and scFv CAR-T against SKOV3, A549, BxPC3, and BxPC3 EGFR knockout (KO) cells co-cultured for 18 h at various E:T ratios followed by a 48 h recovery. **d**, Cytokine and effector protein measurements from supernatants of DNB and scFv CAR-T co-cultured with BxPC3 tumor cells for 18 h. The dotted line represents the assay’s lower limit of detection. **e**, Representative images from live-cell microscopy of co-cultures of T cells transduced with CAR-mCherry fusion proteins and K562 cells transduced with EGFR-eGFP fusion protein, showing immunological synapse formation (white triangles). Raw images shown in **Supplementary Data**. **f**, Growth curves of DNB and scFv CAR-T under weekly re-stimulation (arrows) with irradiated K562 cells expressing EGFR. **g**, Timeline for patient-derived organoid (PDO) and CAR-T co-cultures. Effector CAR T-cells were added at a 10:1 E:T ratio. After 48h of co-culture, live cell imaging and immunoassays were conducted. **h**, Representative images from live-cell microscopy of pancreatic cancer PDOs (specimen B678) co-cultured for 48h with vehicle control, UTD, DNB CAR-T and scFv CAR-T. DNB and scFv CAR-T are marked with a mCherry fluorescent reporter. The experiment was performed using two pancreatic cancer PDO lines (specimen B536 in **Extended Data** Fig. 3b) in technical duplicates with similar results. **i**, Selected cytokines and effector proteins measured in supernatants of DNB and scFv CAR-T co-cultured with pancreatic cancer PDOs (specimen B678) for 48 h (additional analytes are depicted in **Extended Data** Fig. 3c). Data in **b**, **c**, **d**, **f**, **i** are shown as mean ± s.e.m.; n = 3 independent replicates using primary T cells from three donors, each tested in technical triplicates (**b**, **c**) or duplicates (**d**, **f**, **i**); Statistical analysis was performed using a two-way ANOVA (**c** and **f**) or a ordinary one-way ANOVA (**d** and **i)** with Tukey’s multiple comparisons test; P-values indicate the comparison between DNB and scFv CAR-T.

### Comparing DNB- and scFv-based CAR-T

We hypothesized that DNBs would also function as an antigen-binding domain of a CAR and tested the same DNB- and scFv-based EGFR binders in a second-generation CAR construct with otherwise identical transmembrane, 4-1BB co-stimulatory, and CD3zeta signaling domains (**Extended Data Fig. 2a**). We transduced these CARs into primary human T cells and first measured their cytotoxic potency against human EGFR-expressing pancreatic, lung, and ovarian cancer cell lines. DNB-based CAR-T lysed all target cells at levels similar to or higher than the scFv-based CAR-T, despite targeting different sites within EGFR (**Fig. 1c** and **Extended Data Fig. 2b**). In addition, we knocked out (KO) *EGFR* in the BxPC3 cell line using *Streptococcus pyogenes* Cas9 (see **Methods** and **Supplementary Table 2**)^26^. The lack of *EGFR* expression was confirmed by flow cytometry analysis (**Extended Data Fig. 2c**). Notably, lysis of the cells lacking *EGFR* expression was substantially reduced for both DNB- and scFv-based CAR-T. For DNB-based CAR-T, the residual lytic activity was similar to untransduced control T cells (UTDs), consistent with antigen-specific cytotoxicity (**Fig. 1c**). Next, we analyzed supernatants from co-cultures of CAR-T with EGFR-expressing target cells. We found that most cytokines and cytotoxic effector proteins were produced at similar or higher rates by EGFR-targeting DNB CAR-T compared to scFv CAR-T (**Fig. 1d** and **Extended Data Fig. 2d**). Interestingly, among the cytokines secreted at higher levels by DNB-based CAR-T were both pro-inflammatory and anti-inflammatory ones (TNFalpha, IL-2, IL-6, IL-4, IL-10), indicating a robust CAR-T activation and a balanced effector cell response (**Fig. 1d** and **Extended Data Fig. 2d**). To visualize how DNB-based CAR-T engage with target cells, we used fluorescence microscopy (**Fig. 1e**). In co-cultures of T cells expressing CAR-mCherry fusion proteins and K562 scaffold cells expressing an EGFR-GFP fusion protein, co-localization at the CAR-T/target cell interface was observed for both DNB and scFv-based CAR-T, indicating the formation of an immunological synapse (**Fig. 1e**). Next, we assessed the binding avidity of the immune synapse by acoustic force microscopy (AFM). Interestingly, the DNB-based CAR-T displayed robust binding, albeit they had a faster dissociation rate than the scFv CAR-T, consistent with known differences in overall binding strength^7^ (**Extended Data Fig. 2e**). To assess the capability of DNB-based CAR-T to persist and proliferate over an extended period, we tested them in repetitive stimulation assays side-by-side with scFv CAR-T. Both were stimulated weekly with irradiated K562 scaffold cells expressing EGFR, and population doublings were monitored. For the first two weeks, both CAR-T populations grew logarithmically at comparable rates. In the third and fourth weeks, both tapered, albeit this effect was more pronounced for the DNB-based CAR-T (*p* = 0.0003 and <0.0001, respectively) (**Fig. 1f**). Overall, DNB-based CAR-T showed proliferation and persistence upon target antigen encounter, the property most closely associated with beneficial CAR-T therapy outcomes in patients^27–29^. To characterize DNB-based CAR-T in a more clinically relevant model system, we tested their anti-tumor potency against primary patient-derived organoids (PDOs). Pancreatic cancer PDOs that endogenously express EGFR were grown from previously established organoid lines (specimens B536 and B678) (**Fig. 1g** and **Supplementary Table 3**). The organoid lines were cultured for five days to allow for assembly into PDOs before DNB- and scFV-based CAR-T cells were added to the culture. PDO lysis was monitored by confocal microscopy, and at the endpoint, supernatants from 48h co-cultures were harvested for cytokine and effector molecule measurement (**Fig. 1g**). In the control conditions (no treatment and application of untransduced T cells), PDOs maintained their cystic morphology, indicating an intact structure. However, upon treatment with both DNB- and scFv-guided CAR-T cells (both co-expressing mCherry), we observed a gradual infiltration and lysis of the PDOs (**Fig. 1h** and **Extended Data Fig. 3a**). In line with the imaging results, we measured potent release of cytokines (IL-2, TNFα, IFNγ) and Granzyme B from the CAR-T co-cultures with similar or higher levels being secreted from DNB-based compared to scFv-based CAR-T (**Fig. 1i** and **Extended Data Fig. 3b**). In sum, we demonstrate that DNB-guided CAR-T targeting a clinically relevant antigen (EGFR) show similar activity for most short- and long-term T cell effector functions compared to conventional scFv-guided CAR-T.

### RFdiffusion-based BCMA binders in CAR-T

While we were investigating the feasibility of DNB-based CAR-T cell applications, RFdiffusion was introduced as an accessible and easy-to-use generative model for DNB design^9^. We used this model to generate DNBs targeting a new, clinically relevant target: the B cell maturation antigen (in short, BCMA or CD269), also known as “tumor necrosis factor receptor superfamily member 17” (gene name: *TNFRSF17*). Given the small size of the BCMA extracellular domain (54 AA), it was provided in its entirety as the target region for RFdiffusion (PDB identifier 1XU2, chain D)^30^. We used a binder length of 62 AA and a triple-helix bundle as the tertiary structure (**Methods**) (**Fig. 2a**). When running RFdiffusion^9^, we used previously established in silico benchmarks, such as the predicted aligned error (pAE), the predicted local distance difference test (pLDDT), and the root mean-squared deviation (r.m.s.d.) for curating DNB designs (**Methods**). We selected 18 RFdiffusion-generated DNBs predicted to target BCMA for initial experimental testing (**Fig. 2a**; **Extended Data Fig. 4a**; **Supplementary Table 4**). The AI-derived DNBs were incorporated into a second-generation CAR backbone (same as shown in **Extended Data Fig. 2a**), and the resulting constructs were named B1-18 aiCAR. For comparison with an scFv-based CAR, we used a design derived from the FDA-approved idecabtagene vicleucel product (brand name Abecma)^31^. All BCMA-targeting CARs employed identical hinge, transmembrane, and intracellular domains and only differed in their antigen-binding domain. We lentivirally expressed all aiCARs and the control scFv CAR in primary human T cells and screened for functional activity, assessing cytotoxicity and proliferative capacity (**Extended Data Fig. 4b-c**). We selected three of the 18 BCMA-targeting aiCAR designs (B6, B10, B16) that performed favorably in both readouts, indicating short-term and long-term functionality, and proceeded to an in-depth characterization. We measured the aiCAR-T cells’ cytotoxic potency against a panel of human multiple myeloma cell lines that endogenously express BCMA at different levels (**Fig. 2b** and **Extended Data Fig. 4d**). Across all target cells tested, a similar pattern was observed: specific lysis with the scFv-based CAR-T was highest, closely followed by B10 aiCAR-T, and successively decreasing efficiencies with B6 and B16 aiCAR-T (**Fig. 2b**).

These differences were less pronounced for the myeloma cells with high BCMA expression levels (MM.1S and RPMI-8226) and stronger for the cells with lower BCMA expression (L-363), possibly indicating that BCMA-targeting aiCAR-T require higher antigen expression levels for activation (**Fig. 2b** and **Extended Data Fig. 4d**). The trend observed for the different lytic activities was matched by the levels of effector proteins and cytokines secreted from co-cultures of CAR-T and RPMI-8226 myeloma cells (**Fig. 2c**). ScFv-based CAR-T and B10 aiCAR-T secreted Granzyme A, Granzyme B, Perforine, and soluble Fas ligand at comparable levels, while B6 aiCAR-T and B16 aiCAR-T showed successively decreasing levels (**Fig. 2c**). Overall, cytokine secretion was lower for the aiCAR-T compared to the scFv-based CAR-T, with the B10 aiCAR secreting higher levels than the B6 aiCAR and B16 aiCAR (**Fig. 2c**). To confirm long-term effector function, we tested the three main candidates’ capability to persist and proliferate by setting up repetitive stimulation assays. AiCAR-T and scFv CAR-T were stimulated weekly with irradiated K562 scaffold cells expressing BCMA, and CAR-T population doublings were recorded. Over the four-week assay period, B10 aiCAR-T and scFv CAR-T grew at the same rate, with a logarithmic increase in the first three weeks and tapering in the fourth week. Similar to the previously assessed effector functions, B6 aiCAR-T and B16 aiCAR-T showed lower antigen-specific proliferation in comparison to B10 aiCAR-T (**Fig. 2d**). In sum, BCMA-directed aiCARs B6 and B16 were not optimally functional against BCMA-expressing target cells. Therefore, we focused subsequent studies on a closer examination of the B10 aiCAR. First, we measured the binding avidity of B10 aiCAR-T to BCMA-expressing target cells. As previously observed for EGFR-targeting CAR-T, the B10 aiCAR-T displayed robust binding albeit had a faster dissociation rate than the scFv CAR-T (**Extended Data Fig. 4e**).

**Fig. 2.**
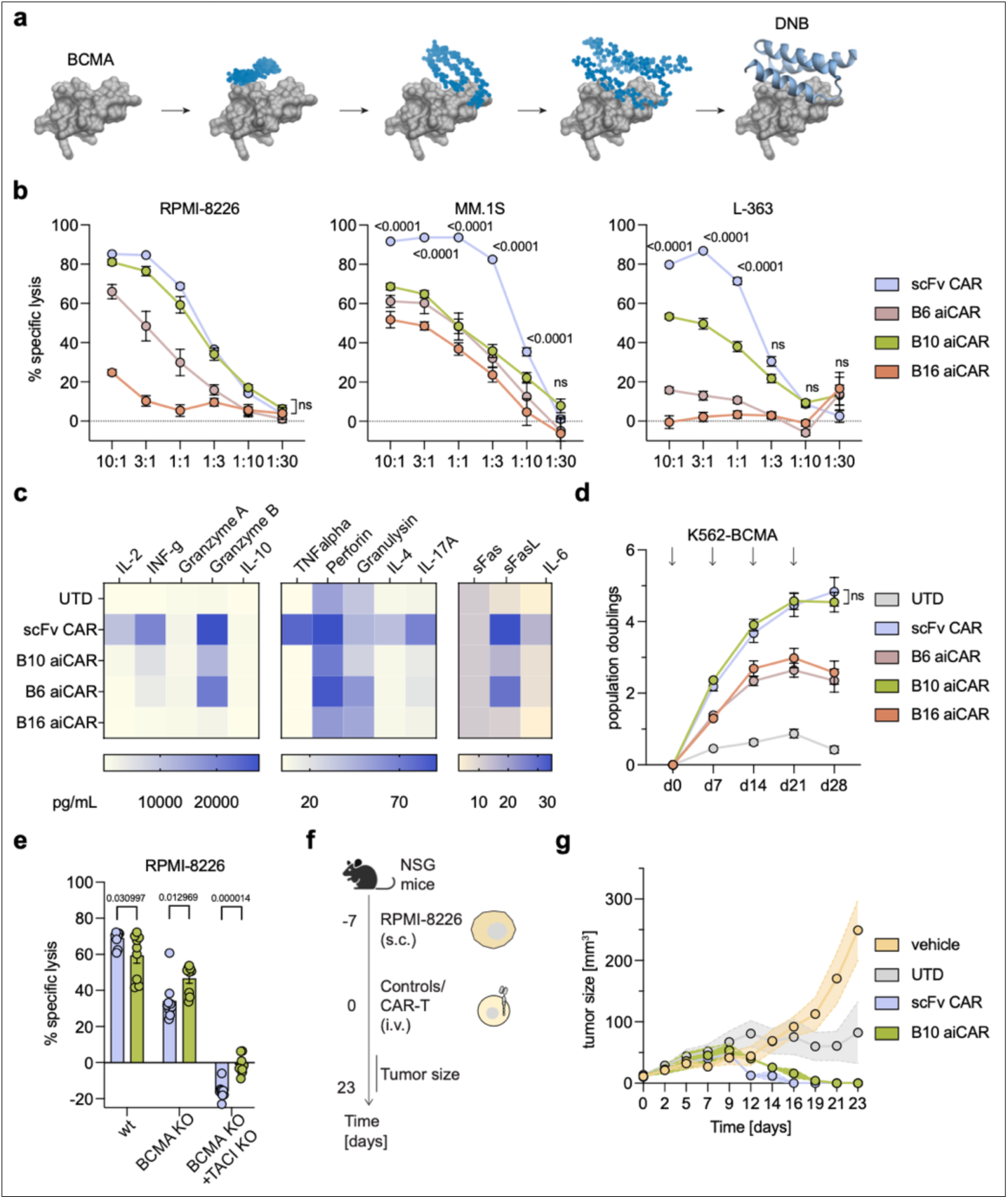
RF diffusion-based binder design for aiCAR-T cells targeting BCMA. **a**, Schematic of the RFdiffusion-based forward denoising process used to generate binders (blue) that target the extracellular domain of BCMA (gray). **b**, Cytotoxicity of ai- and scFv CAR-T against RPMI-8226, MM.1S, and L-363 co-cultured for 16h at various E:T ratios and normalized to the UTD control. **c**, Heatmap of mean secreted cytokine and effector proteins from 16h co-cultures of ai- and scFv CAR-T with RPMI-8226 tumor cells (1:1 E:T ratio). **d**, Growth curves of UTD controls, ai- and scFv CAR-T under weekly re-stimulation (arrows) with irradiated K562 cells expressing BCMA. **e**, Cytotoxicity of B10 aiCAR-Ts and scFv CAR-T against RPMI-8226, RPMI-8226 with TNFRSF17 (BCMA) KO, and RPMI-8226 with TNFRSF17 and TNFSF13B (TACI) double-KO after 16h co-culture at a 1:1 E:T ratio. **f**, Schematic of RPMI-8226 in vivo model treated with 5x10^6^ CAR-T intravenously 7 days after subcutaneous engraftment of 5x10^6^ RPMI-8226 cells. Tumor burden was monitored by caliper measurement over time. **g**, Tumor volume curves of the four treatment groups (vehicle, UTD, scFv- und B10 aiCAR-T) (n = 5 mice per condition). Data in **b**, **d**, **e**, **g** are shown as mean ± s.e.m. Data in **b-e** were obtained from n = 3 donors, each in technical triplicates (**b**, **e**) or duplicates (**c**, **d**). Statistical analysis was performed using two-way ANOVA with Tukey’s multiple comparisons tests (**b** and **d**) or multiple unpaired t-tests (**e**); P-values indicate the comparison between aiCAR-T and scFv CAR-T.

Next, we aimed to assess the specificity of B10 aiCAR-T by testing them in RPMI-8226 myeloma cells with a *TNFRSF17* KO, i.e. in cells lacking BCMA expression (**Methods** and **Extended Data Fig. 4d**). Unexpectedly, we observed that the KO of *TNFRSF17 (*the gene encoding BCMA) did not abrogate lytic activities in either scFv- or in B10-based CAR-Ts. However, lytic activities were reduced from 68% and 59% in wild-type (WT) cells (scFV vs. AI-CAR; means of specific lysis; *p=0.03*) to 34% and 47% (*p=0.013)* in KO cells, respectively (**Fig. 2e**). As a potential explanation for why the lytic activity of B10 aiCAR was less affected by the KO of *TNFRSF17* than the activity of the scFv-CAR, we hypothesized that our binder may be demonstrating bispecific binding. For the RFdiffusion input that yielded binder B10, we used a structure of BCMA in complex with its natural ligand, “a proliferation-inducing ligand” (APRIL, CD256; gene name: *TNFSF13*) (PDB identifier 1XU2)^30^. Since APRIL binds to both BCMA and the “transmembrane activator and CAML interactor” (TACI, CD267; gene name: *TNFSF13B*), a member of the tumor necrosis factor receptor superfamily that is closely related to BCMA, we hypothesized that the B10 aiCAR may mimic APRIL’s binding, leading to TACI cross-reactivity. To investigate this, we measured the cytotoxic activity of B10 aiCAR-T against RPMI-8226 myeloma cells in which both *TNFRSF17 and TNFSF13B* were knocked out (BCMA KO, TACI KO)(**Methods, Extended Data Fig. 4d**, and **Fig. 2e**).

As predicted, cells with a double-KO completely abrogated the lytic activity of both B10 aiCAR-T and of scFv-based CAR-T (**Fig. 2e**). This finding indicates a dual specificity against the two target antigens, BCMA and TACI, for our RFdiffusion-generated B10 aiCAR.

### Application of AI-based CAR-T in vivo

To characterize B10 aiCAR-T in a more complex model system, we assessed its anti-tumor potency in a xenograft mouse model of multiple myeloma, side-by-side with the scFv-based anti-BCMA CAR-T (**Fig. 2f**). NSG mice were subcutaneously injected with RPMI-8226 myeloma cells and tumor engraftment was confirmed via caliper measurement before initiation of treatment. Mice received a single intravenous injection of vehicle, UTD, scFv CAR-T, or B10 aiCAR-T. Tumor volumes were monitored three times a week by caliper measurement, and body weight was recorded (**Extended Data Fig. 5a**). Tumor volumes rapidly progressed in the vehicle-treated group and more slowly increased in the mice that had received UTD, indicating some allogeneic rejection, which is often observed in multiple myeloma models. In contrast, scFv CAR-T and B10 aiCAR-T cleared the myeloma tumors in all mice over time (**Fig. 2f-g**). Interestingly, the tumor clearance took 5 days longer (d21 vs. d16) for the B10 aiCAR-T than the scFv CAR-T (**Fig. 2g** and **Extended Data Fig. 5b; Supplementary Table 5)**. This may reflect a lower activation level of B10 aiCAR-T in comparison to the scFv CAR-T, which would align with the previously observed reduction in lytic function and cytokine secretion in vitro.

### Using aiCAR-T to counter cancer resistance

Currently, three classes of BCMA-targeted therapies are in clinical use for relapsed/refractory multiple myeloma patients, either as single agents or combination therapy: CAR-T, TCE, and antibody-drug conjugates. BCMA-targeted therapies show high response rates, but responses are of limited duration, with the majority of patients relapsing within a year^32,33^. Genomic deletions and substitutions in the extracellular domain of *TNFRSF17* have been described as a prevalent mechanism of antigen escape in response to BCMA-targeting therapies^34–36^. Notably, such mutations that render the antigen unrecognizable by the BCMA-targeting therapy occur more frequently in response to TCE and antibody drug conjugates than CAR-T^37^. We hypothesized that we could target such mutant forms of BCMA with our newly generated aiCAR-T cells. A recently approved cellular immunotherapy for relapsed or refractory myeloma is teclistamab, a T-cell redirecting bispecific antibody binding both BCMA and CD3^38^. In myeloma patients who relapsed after receiving teclistamab, the *TNFRSF17* resistance variant encoding BCMA-R27P was recently identified (**Fig. 3a**)^36^. Based on the predicted binding site of B10 to BCMA, we assumed that B10 aiCAR-T could still bind and kill cells expressing the BCMA-R27P variant (**Fig. 3a**). To test our hypothesis, we transduced K562 cells to express either unmodified (WT) BCMA or BCMA-R27P (**Extended Data Fig. 6a**). We first confirmed that the BCMA-R27P variant mediates resistance to T cell killing in the presence of teclistamab in a dose-dependent manner (**Fig. 3b**). Next, we assessed co-cultures of B10 aiCAR-T against K562-BCMA (WT) and K562-BCMA-R27P cells and observed that B10 aiCAR-T cells lysed significantly more K562 cells expressing any form of BCMA than control cells lacking BCMA expression (p<0.0001). Most importantly, cells expressing the BCMA-R27P variant were lysed at similar frequencies to cells expressing WT BCMA (K562-BCMA) by B10 aiCAR-T (*p=0.5556*) (**Fig. 3c** and Extended Data Fig. 6b).

**Fig. 3.**
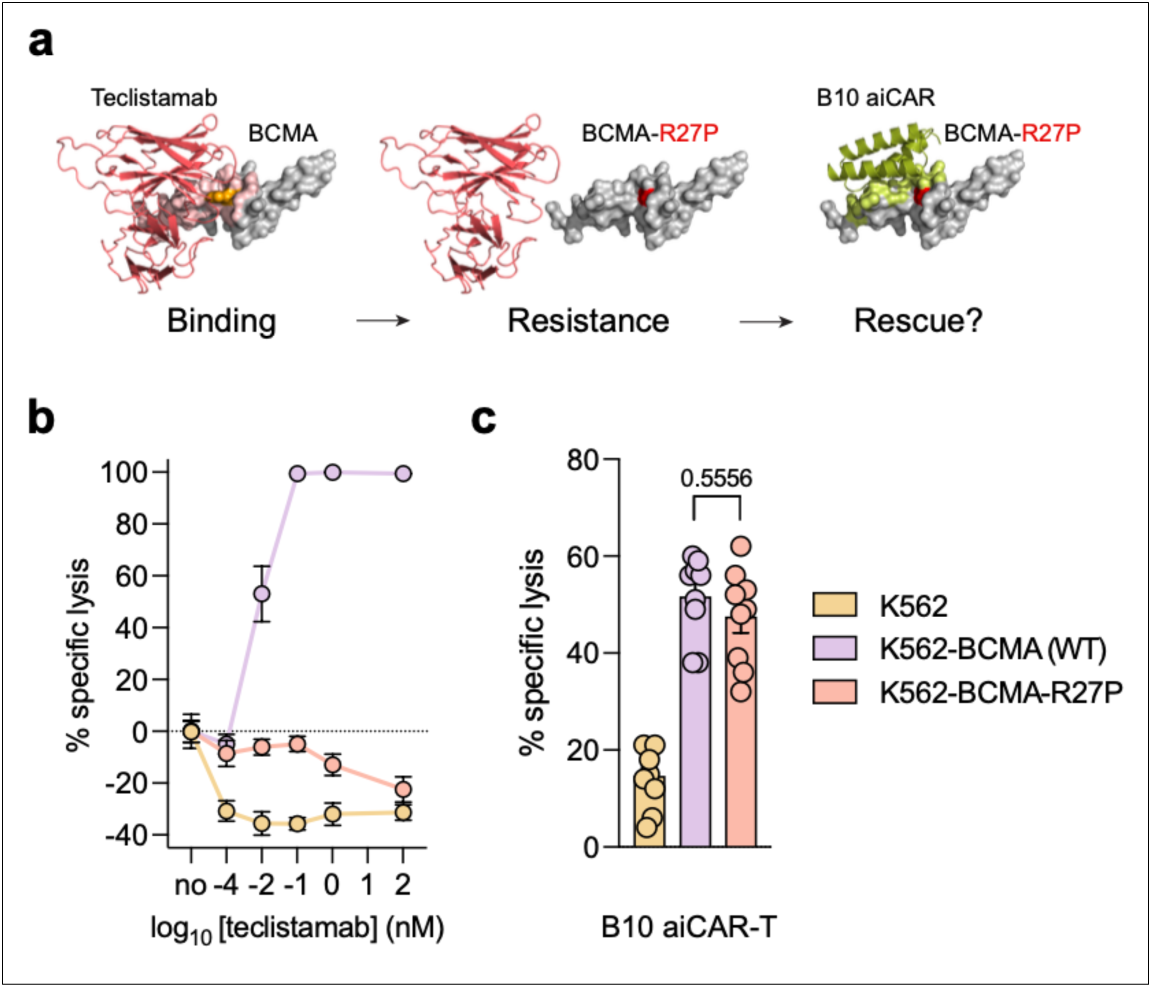
Extending aiCAR-T to tumor antigen resistance variants. **a**, Schematic of the BCMA surface mutation R27P (R27:orange, P27: red) interrupting binding of teclistamab (salmon, with predicted 5Å binding interface in light salmon) and the predicted binding site of B10 (green, with predicted 5Å binding interface in light green) on the extracellular domain of BCMA (gray). **b**, T cell-mediated lysis of K562, K562-BCMA (WT), and K562-BCMA-R27P cells in the presence of various concentrations of teclistamab. Lysis was measured by flow cytometry and normalized to the UTD control. **c**, B10 aiCAR-T specific lysis of K562, K562-BCMA (WT), and K562-BCMA-R27P at a 1:1 E:T ratio. Data in **b**, **c** are shown as mean ± s.e.m.; n = 3 donors, each in technical triplicates; Statistical analysis was performed using an ordinary one-way ANOVA with Tukey’s multiple comparisons test; P-values indicate the comparison between aiCAR-T and scFv CAR-T.

## Discussion

While CAR-T have been successfully tested using a wide array of binding moieties, these domains are often slow and expensive to generate and require methods inaccessible to most laboratories. Our findings serve as proof of concept that in silico-generated DNBs can be used for CAR-T therapies, including self-made binders using a diffusion model. We also demonstrate that aiCAR-T can be used to target a mutated epitope that has evolved to resist a clinically used immunotherapy.

While our data indicate that DNB-based CAR-T can have similar functional properties as scFv-based ones, avidity was reduced in our measurements, and in vivo efficacy will have to be characterized in more detail across different targets and model systems in the future. One aspect for further improvement of aiCAR-T includes “in silico maturation” of DNBs using partial diffusion, potentially enhancing affinity^39^ as well as new methods using backpropagation to hallucinate binders^40^. Moreover, it will be crucial to assess aspects of DNB specificity and safety. Both the potential for off-target binding and immunogenicity using AI-derived DNBs will need to be addressed in future studies. With regard to the targeting of cancer resistance, it would be important to model resistance to aiCAR-T and to use generative AI to design bespoke, mutation-specific binders. This approach would allow highly personalized responses to cancer resistance in a much-accelerated fashion. Of note, the structure-based approaches used here, i.e., RFdiffusion and AF2, are somewhat constrained by using a “static”, single-conformation protein as input. In the near future, the prediction of alternative protein conformations^41^ may alleviate this limitation, enabling DNB designs that take into account different conformations of the same protein, thereby improving binder functionalities. Alternatively, protein language models (pLMs)^42^ have been adapted to generate DNBs based on target sequence alone, without requiring target structures^43^.

In sum, we envision AI-guided DNBs to democratize and accelerate the design of new and bespoke CAR-T therapies. This is important specifically in the context of cancer resistance, which affects most patients receiving CAR-T treatments^6,44^.

Much additional research is needed to better understand and engineer efficacious and safe DNBs. However, aiCAR-T substantially expand the scope of cellular immunotherapy and effectively showcase the application of AI-guided gene & cell therapies.

## Acknowledgments

This work was supported by the German Research Foundation (DFG): Emmy Noether Program to J.G. (no. 502106724) and A.S. (no. 494809850); Start-up funding from TRR 338/1 2021 – 452881907 to A.S.; TRR 387/1 – 514894665, DFG BA 2851/6-1 (project ID: 452409123) and DFG BA 2851/7-1 (project ID: 537477296) to F.B.; TRR 338/1 2021 - 452881907 (project A01) to D.H.B.; M.M. received support from a Translational Medicine Program scholarship from the Technical University of Munich. J.G. was supported by the Aventis Foundation.

We thank Katja Heinrich, Tatjana Nedelko, Julia Höbart, Linda Bachmann, Irene Santisteban Oritz, Monique Pena Schäfer, Elvira D’Ippolito, and Nadine Glaser for technical assistance. We thank Simon T. Schäfer, Sebastian Kobold, Benjamin P. Kleinstiver, Markus Elsner, J. Keith Joung, and Marcela V. Maus for the helpful discussions. We thank David Baker and his laboratory for providing broad access to RFdiffusion. We also thank Sergey Ochinnikov, who provided the Google Collab of RFdiffusion that we used for this work.

## Author contributions

M.M., D.A., J.G., and A.S. designed the project and experiments. M.M., D.A., N.K., A.SF., A.C., and V.L. performed the experiments. M.M., D.A., N.K., A.SF., M.S., N.H., A.C., L.R., and V.L. developed the methods. M.R., K.-L.L., F.B., D.H.B., J.G., and A.S. provided resources and oversight. M.M., D.A., J.G., and A.S. wrote the manuscript receiving input from all authors.

## Competing interests statement

J.G. is a paid consultant for Poseida Therapeutics, a company developing various gene and cell therapies, and has financial interests in the company. J.G. is a member of the gene editing committees of ASGCT and ESGCT. A.S. is a paid consultant for TQ Therapeutics, a company developing gene and cell therapies. M.R. has received payments or honoraria for lectures, presentations, and participation in speakers’ bureaus from Celgene, Falk, Servier, and Roche. F.B. received honoraria and/or travel/accommodation expenses from BMS, AbbVie, and Janssen. M.M., D.A., J.G., and A.S. are co-inventors on a patent application filed by TUM on engineering DNB-based CAR-T therapies.

Correspondence and requests for materials should be addressed to J.G. or A.S..

## Extended Data Figures

**Extended Data Fig. 1.**
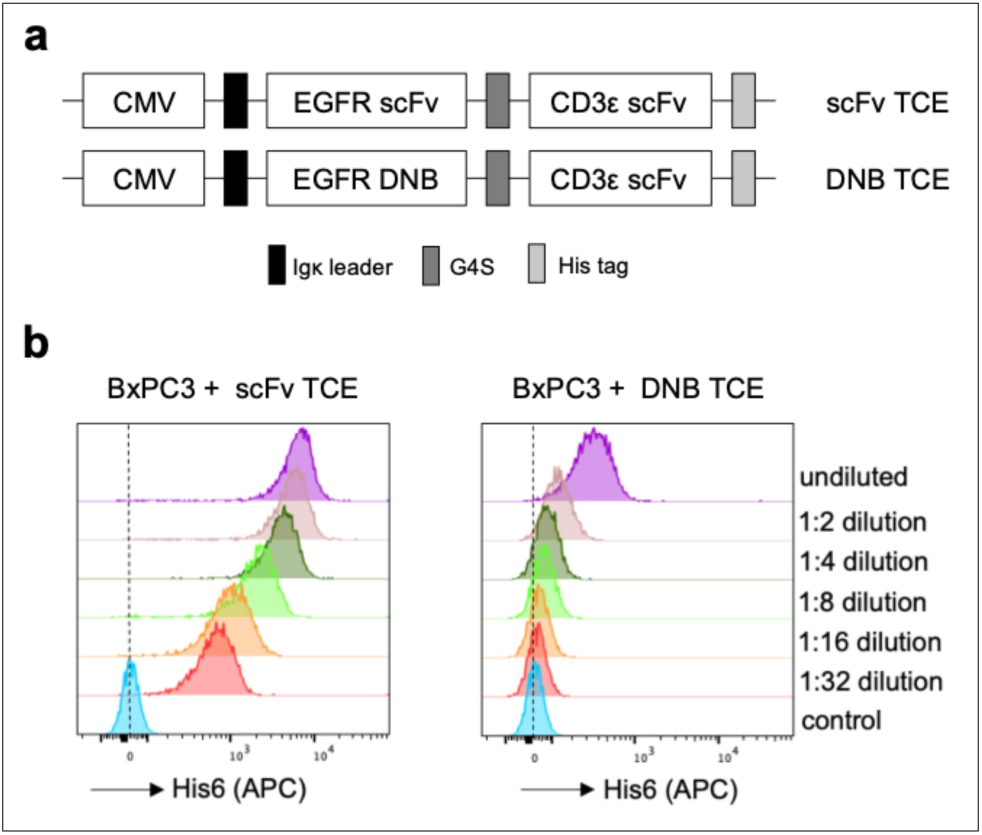
Design of DNB-based TCEs and their binding to EGFR target cells. **a**, Schematic of TCE constructs targeting EGFR. **b,** Flow cytometry histograms demonstrating dose-dependent binding of the indicated TCE to BxPC3 cells, detected via His-tag staining. HEK supernatant from untransduced cells served as a negative control. n = 3 independent replicates; representative histograms are shown.

**Extended Data Fig. 2.**
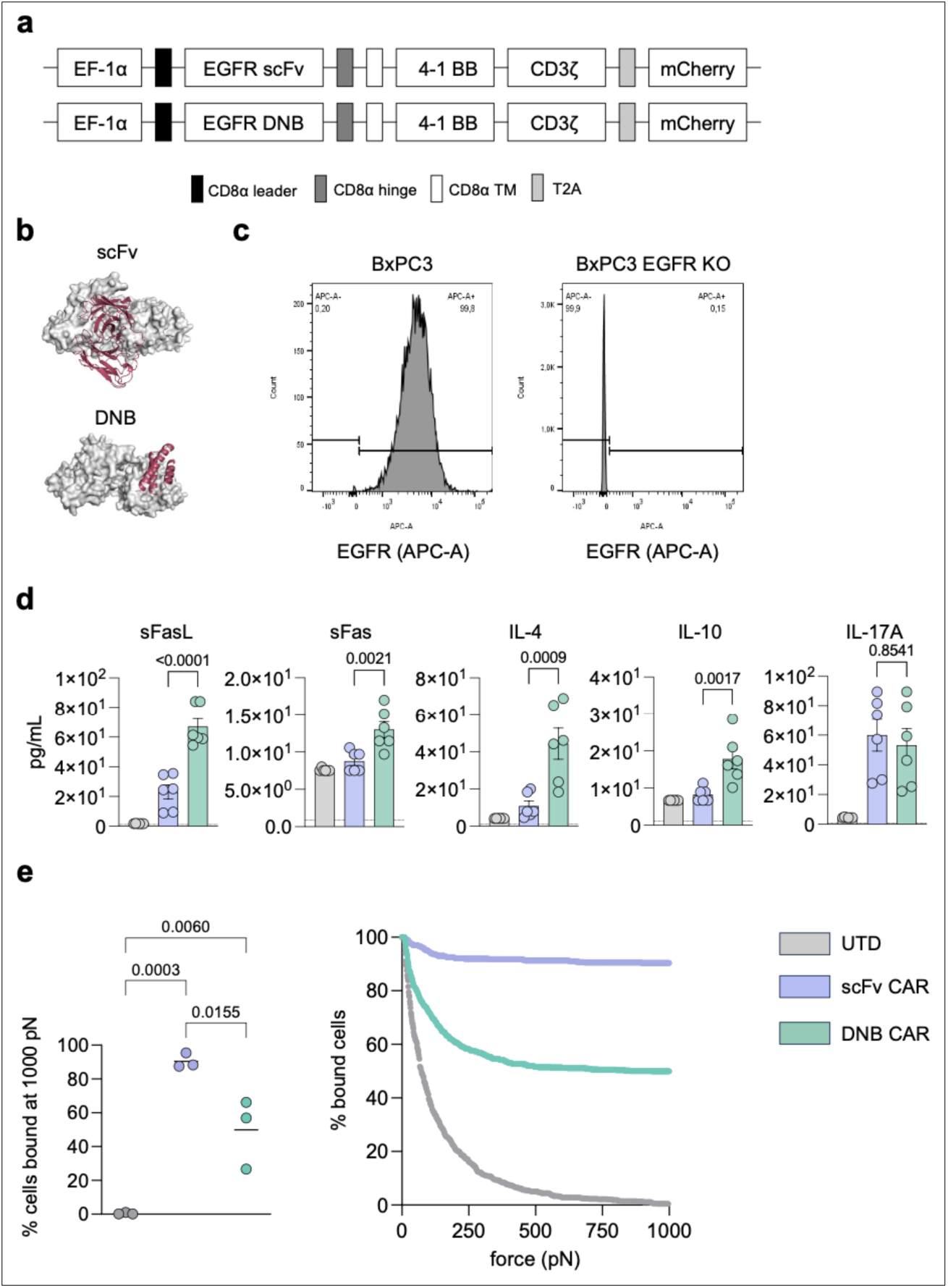
Design of DNB-based CARs and their functional characterization. **a**, Schematic of DNB and scFv CAR-T constructs. **b**, Schematic illustrating the scFv (top) and DNB (bottom) targeting the extracellular domain of EGFR. **c**, EGFR staining of human pancreatic cancer cell lines (BxPC3, BxPC3 with EGFR KO) using conjugated flow antibodies. **d**, Additional cytokine and effector protein measurements from supernatants of DNB and scFv CAR-T co-cultured with BxPC3 tumor cells for 18 h. **e**, Avidity of CAR-Ts on BxPC3 target cells after 15min co-incubation measured by acoustic force microfluidic microscopy (AFMM). The dot plot displays the normalized percentage of bound UTDs (gray), scFv CAR-T (blue), DNB CAR-T (green) on BxPC3 tumor cells at 1000 pN endpoint (left panel). The avidity graph shows the normalized percentage of bound cells with increase in acoustic force of one representative measurement out of three (right panel). Data in **d** are shown as mean ± s.e.m.; n = 3 donors, each in technical duplicates. Data in **e** are shown as mean; n = 3 independent replicates. Statistical analysis was performed using an ordinary one-way ANOVA with Tukey’s multiple comparisons test.

**Extended Data Fig. 3.**
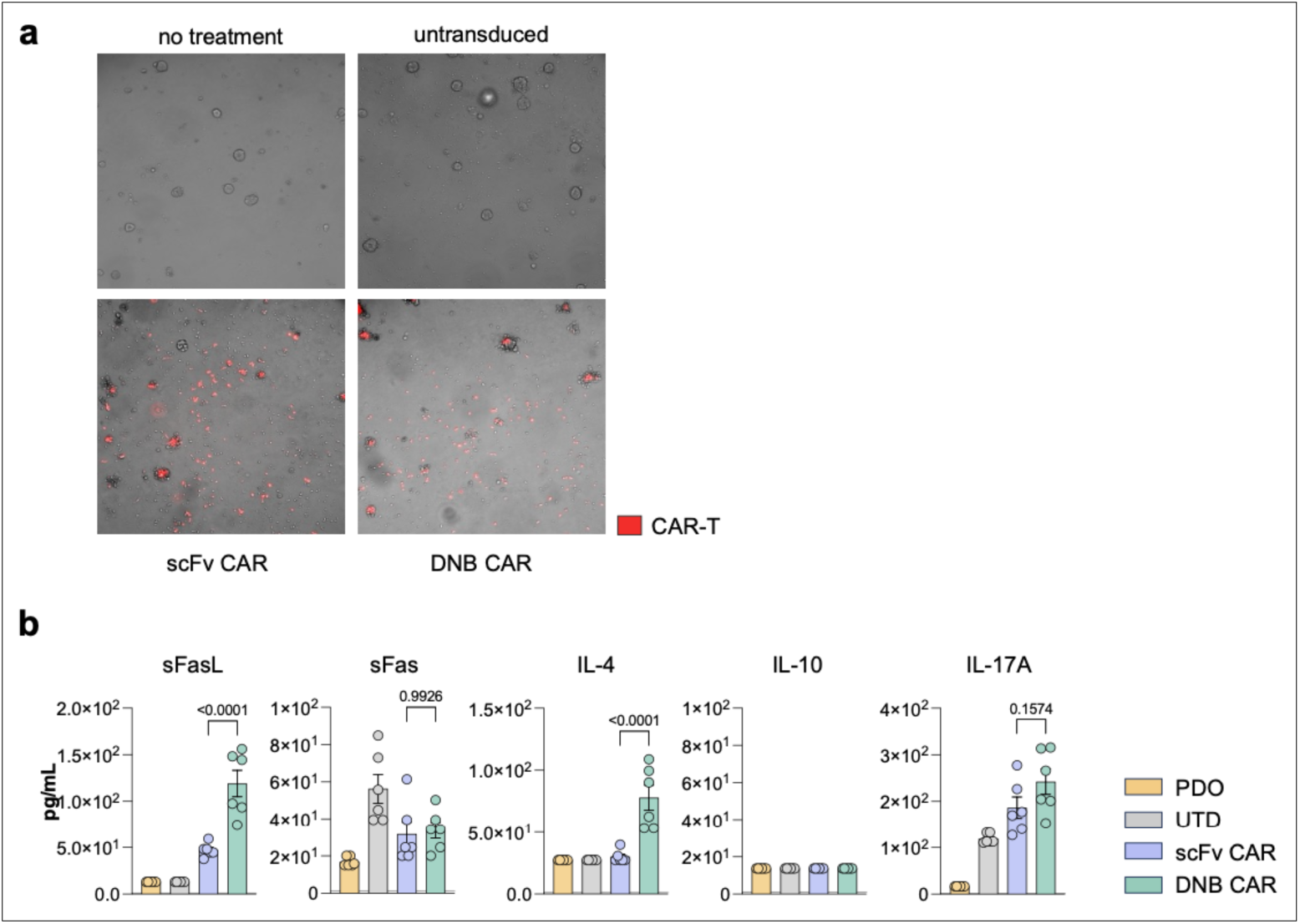
Activity of DNB CAR-T against pancreatic cancer PDOs. **a**, Representative images from live-cell microscopy of pancreatic cancer PDOs (specimen B536) co-cultured for 48h with vehicle control, UTD, DNB CAR-T and scFv CAR-T. DNB and scFv CAR-T are marked with a mCherry fluorescent reporter. **b**, Additional cytokines and effector proteins measured in supernatants of DNB and scFv CAR-T co-cultured with pancreatic cancer PDOs (specimen B678) for 48 h. Data in **b** are shown as mean ± s.e.m.; n = 3 donors, each in technical duplicates. Statistical analysis in **b** was performed using an ordinary one-way ANOVA with Tukey’s multiple comparisons test.

**Extended Data Fig. 4.**
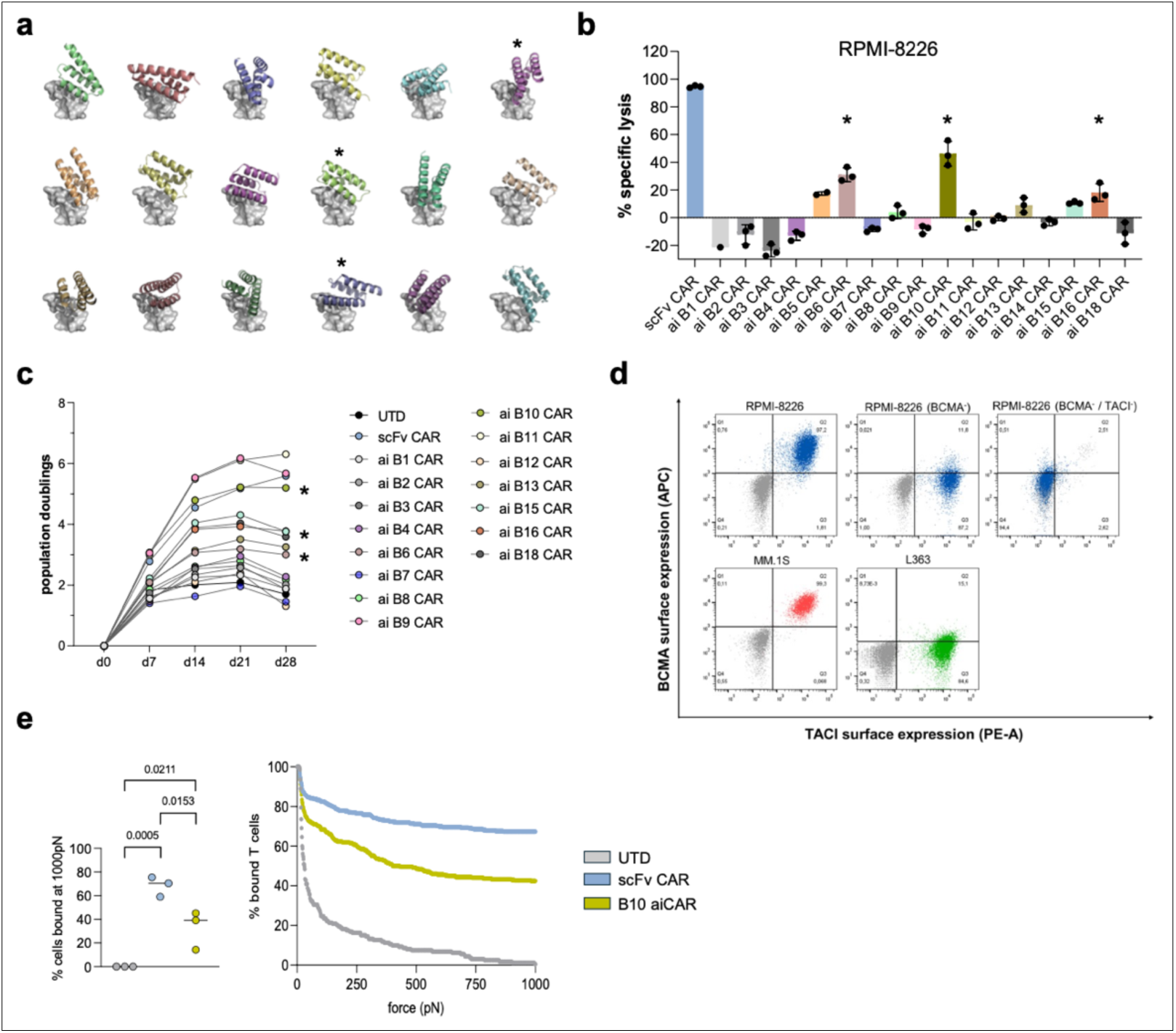
Functional screening of BCMA-targeting aiCAR-T and characterization of lead candidates. **a**, Schematic of the 18 RFdiffusion-generated DNBs targeting BCMA. **b**, Cytotoxicity screen of ai- and scFv CAR-T against RPMI-8226 co-cultured for 16h at 1:1 effector:target ratio and normalized to the UTD control. **c**, Growth curves of UTD, ai- and scFv CAR-T under weekly re-stimulation with irradiated K562 cells expressing BCMA. **d**, BCMA and TACI staining of multiple myeloma cell lines RPMI-8226, RPMI-8226 with TNFRSF17 (BCMA) KO, and RPMI-8226 with TNFRSF17 and TNFSF13B (TACI) double-KO, as well as MM.1S and L363 using conjugated flow antibodies (stained cells in blue, green or red, isotype stain in gray). **e**, B10 aiCAR-T and scFv CAR-T avidity to MM.1S target cells measured by AFMM after a 15 min co-incubation. The dot plot displays the normalized percentage of bound UTDs (gray), scFv CAR-T (blue), B10 aiCAR-T (green) on MM.1S tumor cells at 1000 pN endpoint (left panel). The avidity graph shows the normalized percentage of bound cells with increase in acoustic force of one representative measurement out of three (right panel); n = 3 independent experiments; statistical analysis was performed using one-way ANOVA with Tukey’s multiple comparisons test; P-values indicate the comparison between aiCAR-T and scFv CAR-T. Asterisks in **a**, **b,** and **c** mark the DNBs selected for subsequent characterization. Data in **b** are shown as mean ± s.d. Data in **b**, **c** were obtained from n = 1 donor in technical triplicates (**b**) or duplicates (**c**). Statistical analysis in **e** was performed using an ordinary one-way ANOVA with Tukey’s multiple comparisons test.

**Extended Data Fig. 5.**
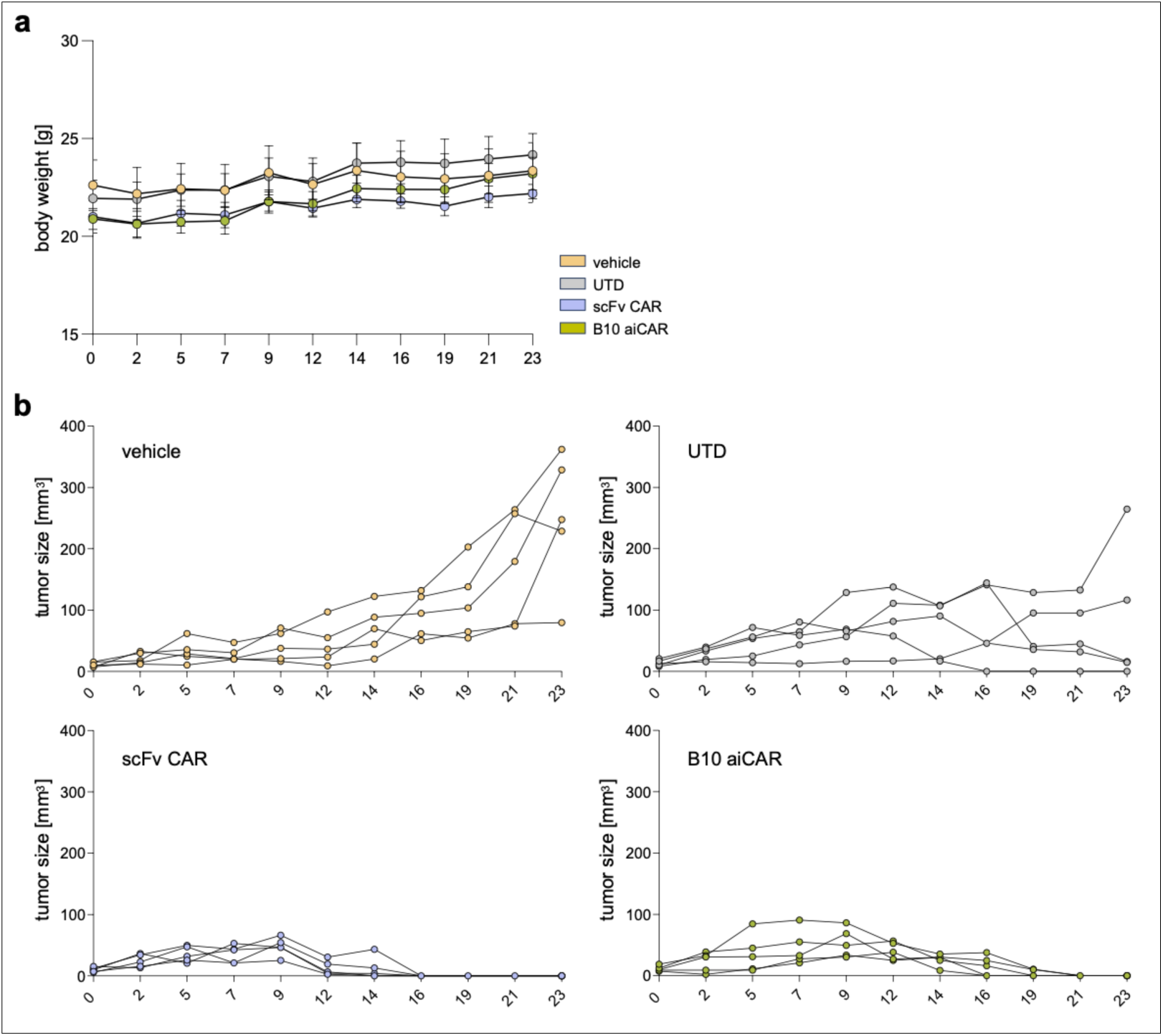
Activity of B10 aiCAR-T in a xenograft mouse model of mulitple myeloma. **a**, Weight curves of the four treatment groups (vehicle, UTD, scFv-und B10 aiCAR-T) (n = 5 mice per condition). **b**, Individual tumor volume curves of the four treatment groups (vehicle, UTD, scFv-und B10 aiCAR-T) (n = 5 mice per condition). Data in **a** and **b** are shown as mean ± s.e.m.

**Extended Data Fig. 6.**
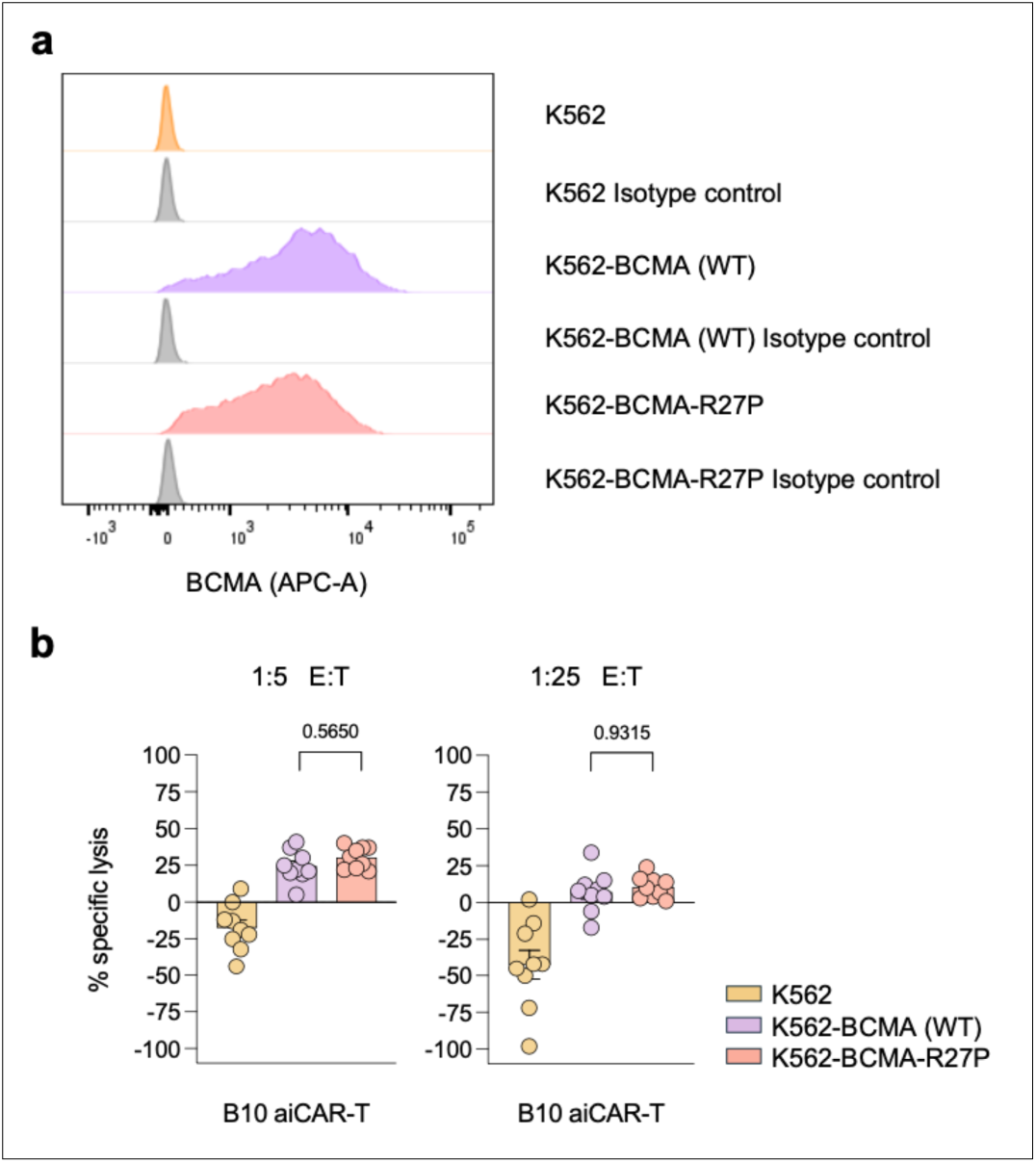
**a**, BCMA surface expression in the generated K562 cell lines using polyclonal anti-BCMA antibodies (colored) or isotype control antibodies (gray) for flow cytometry. **b**, B10 aiCAR-T specific lysis of K562, K562-BCMA (WT), and K562-BCMA-R27P at a 1:5 (left) and 1:25 (right) E:T ratio. Data in **b** are shown as mean ± s.e.m.; n = 3 donors, each in technical triplicates; Statistical analysis was performed using an ordinary one-way ANOVA with Tukey’s multiple comparisons test.

## Supplementary Tables

**Supplementary Table 1:**
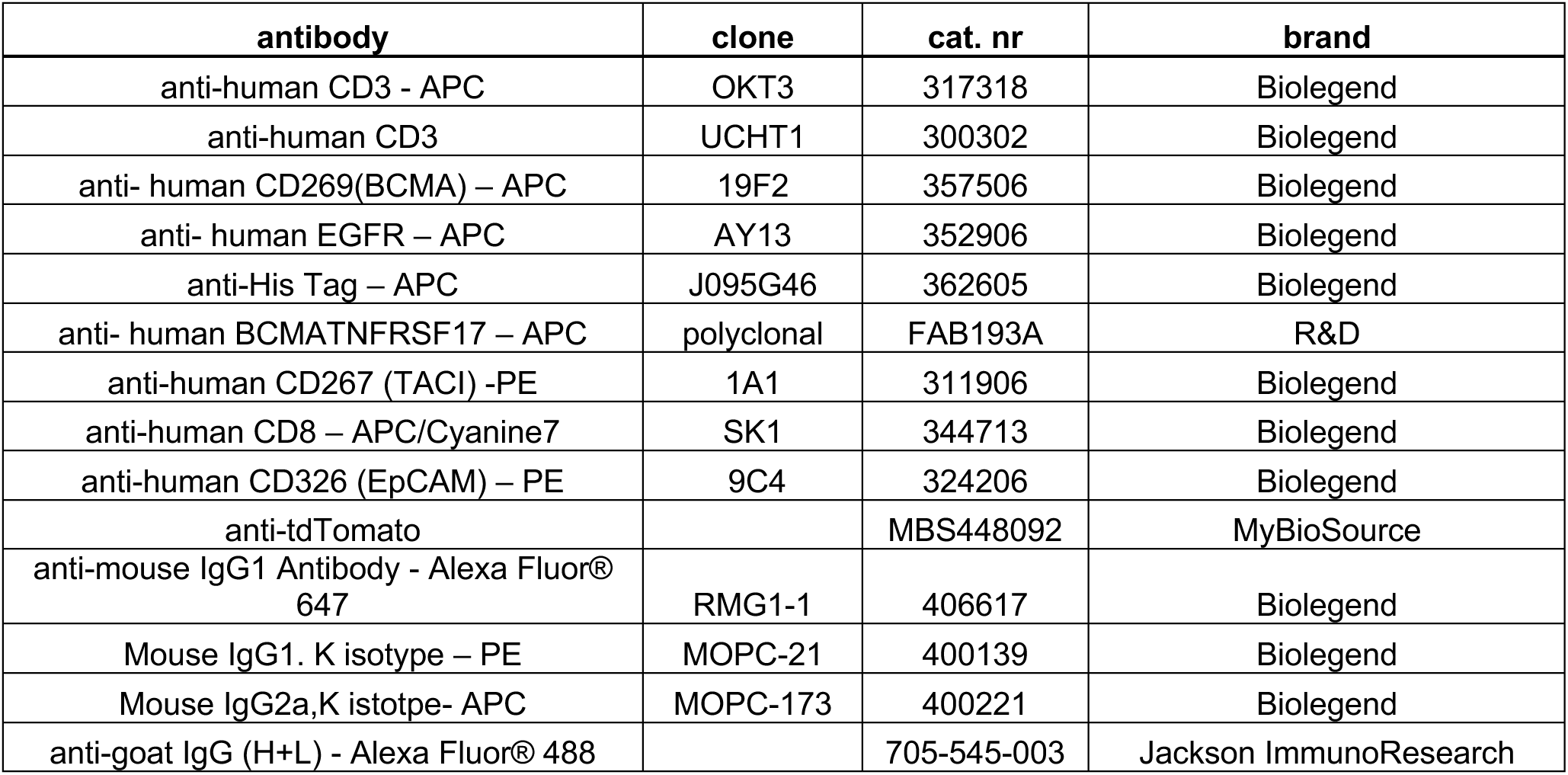
Antibodies.

| antibody | clone | cat. nr | brand |
| --- | --- | --- | --- |
| anti-human CD3 - APC | OKT3 | 317318 | Biolegend |
| anti-human CD3 | UCHT1 | 300302 | Biolegend |
| anti- human CD269(BCMA) – APC | 19F2 | 357506 | Biolegend |
| anti- human EGFR – APC | AY13 | 352906 | Biolegend |
| anti-His Tag – APC | J095G46 | 362605 | Biolegend |
| anti- human BCMATNFRSF17 – APC | polyclonal | FAB193A | R&D |
| anti-human CD267 (TACI) -PE | 1A1 | 311906 | Biolegend |
| anti-human CD8 – APC/Cyanine7 | SK1 | 344713 | Biolegend |
| anti-human CD326 (EpCAM) – PE | 9C4 | 324206 | Biolegend |
| anti-tdTomato |  | MBS448092 | MyBioSource |
| anti-mouse IgG1 Antibody - Alexa Fluor® 647 | RMG1-1 | 406617 | Biolegend |
| Mouse IgG1. K isotype – PE | MOPC-21 | 400139 | Biolegend |
| Mouse IgG2a,K isotpe- APC | MOPC-173 | 400221 | Biolegend |
| anti-goat IgG (H+L) - Alexa Fluor® 488 |  | 705-545-003 | Jackson ImmunoResearch |

**Supplementary Table 2:**
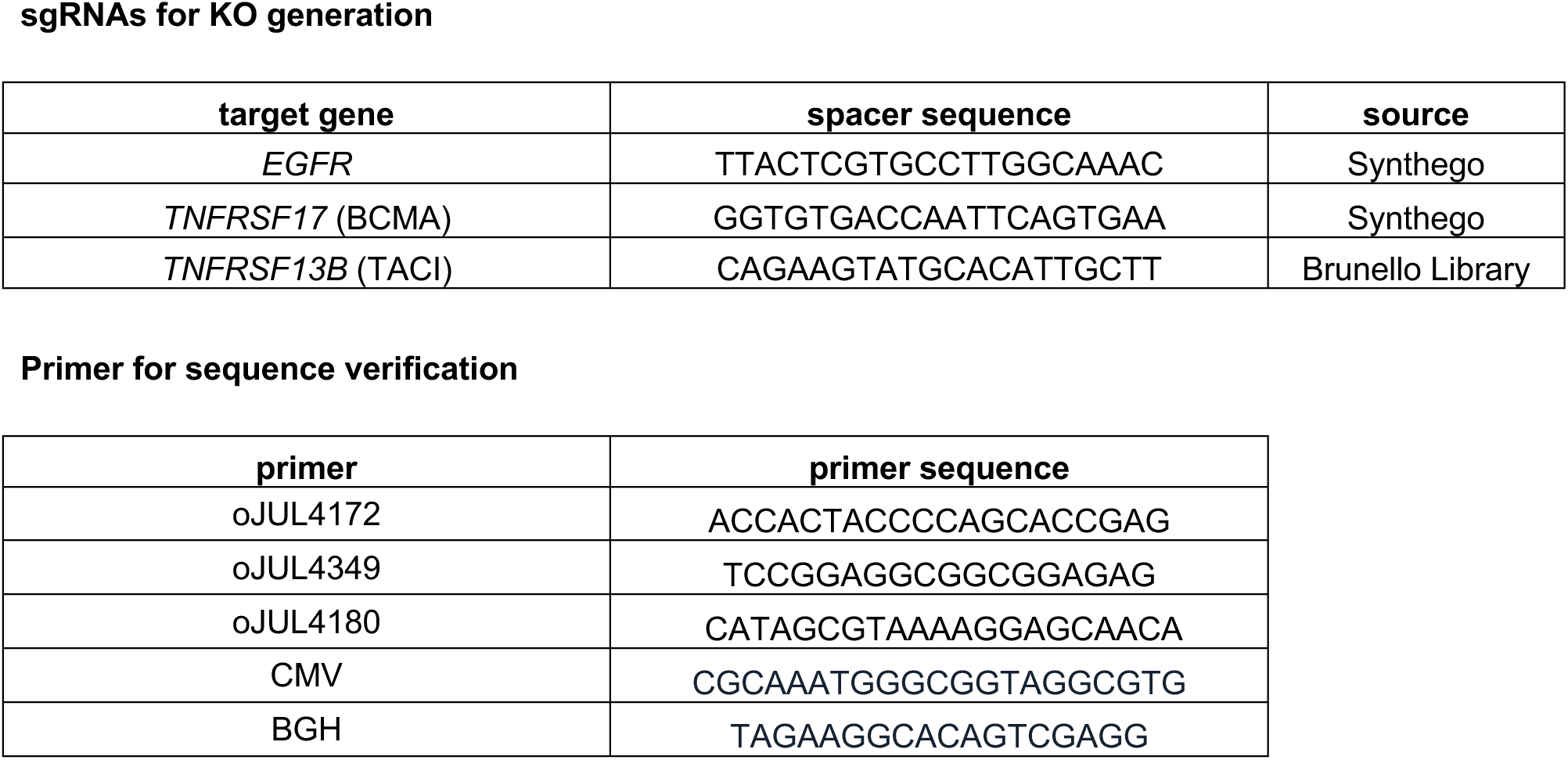
Primer und SpCas9 single guide RNA (sgRNA) spacer.

**Supplementary Table 3:**
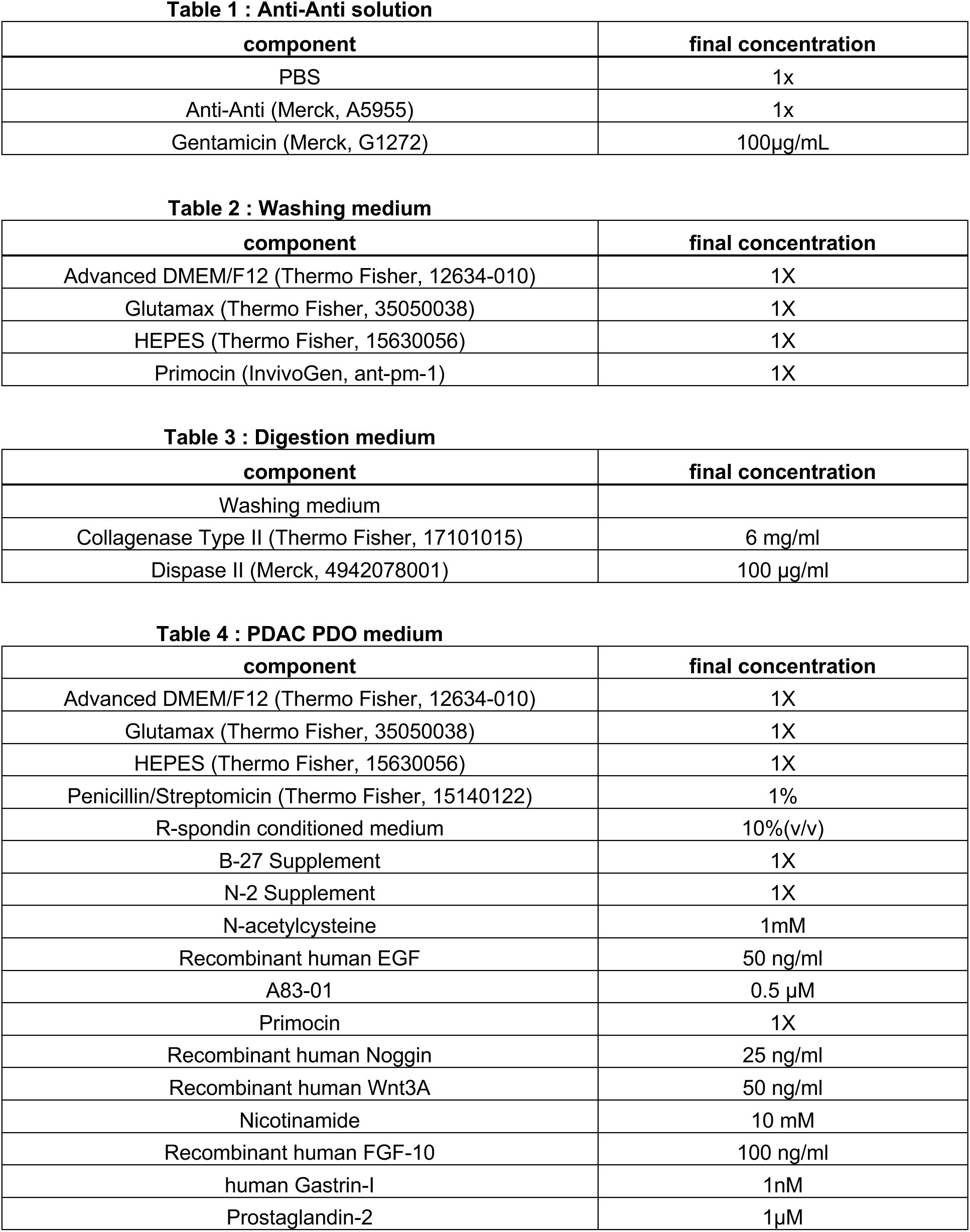
Generation and culture of PDAC patient-derived organoids.

| **component** | **final concentration** |
| --- | --- |
| PBS | 1x |
| Anti-Anti (Merck, A5955) | 1x |
| Gentamicin (Merck, G1272) | 100µg/mL |

| **component** | **final concentration** |
| --- | --- |
| Advanced DMEM/F12 (Thermo Fisher, 12634-010) | 1X |
| Glutamax (Thermo Fisher, 35050038) | 1X |
| HEPES (Thermo Fisher, 15630056) | 1X |
| Primocin (InvivoGen, ant-pm-1) | 1X |

| **component** | **final concentration** |
| --- | --- |
| Washing medium |  |
| Collagenase Type II (Thermo Fisher, 17101015) | 6 mg/ml |
| Dispase II (Merck, 4942078001) | 100 µg/ml |

| **component** | **final concentration** |
| --- | --- |
| Advanced DMEM/F12 (Thermo Fisher, 12634-010) | 1X |
| Glutamax (Thermo Fisher, 35050038) | 1X |
| HEPES (Thermo Fisher, 15630056) | 1X |
| Penicillin/Streptomycin (Thermo Fisher, 15140122) | 1% |
| R-spondin conditioned medium | 10%(v/v) |
| B-27 Supplement | 1X |
| N-2 Supplement | 1X |
| N-acetylcysteine | 1mM |
| Recombinant human EGF | 50 ng/ml |
| A83-01 | 0.5 µM |
| Primocin | 1X |
| Recombinant human Noggin | 25 ng/ml |
| Recombinant human Wnt3A | 50 ng/ml |
| Nicotinamide | 10 mM |
| Recombinant human FGF-10 | 100 ng/ml |
| human Gastrin-I | 1nM |
| Prostaglandin-2 | 1µM |

**Supplementary Table 4:**
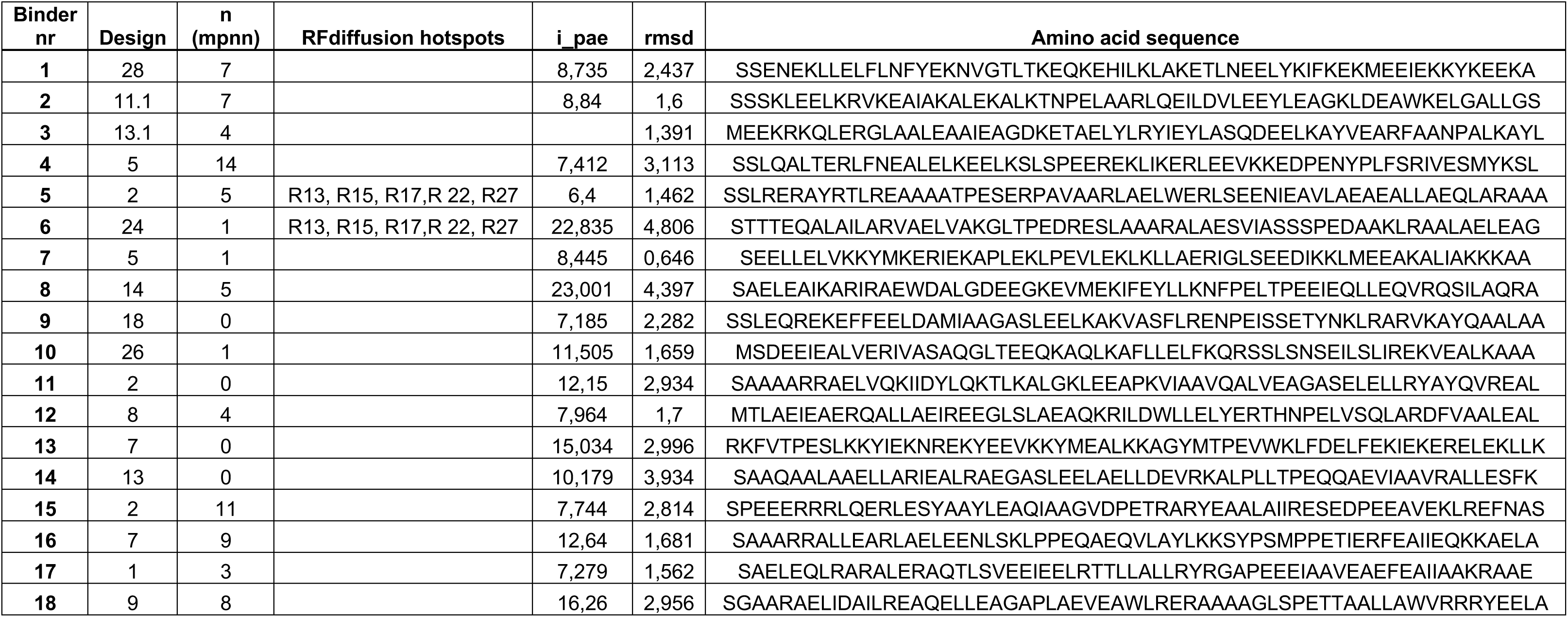
RFdiffusion binder prompts and values.

**Supplementary Table 5:**
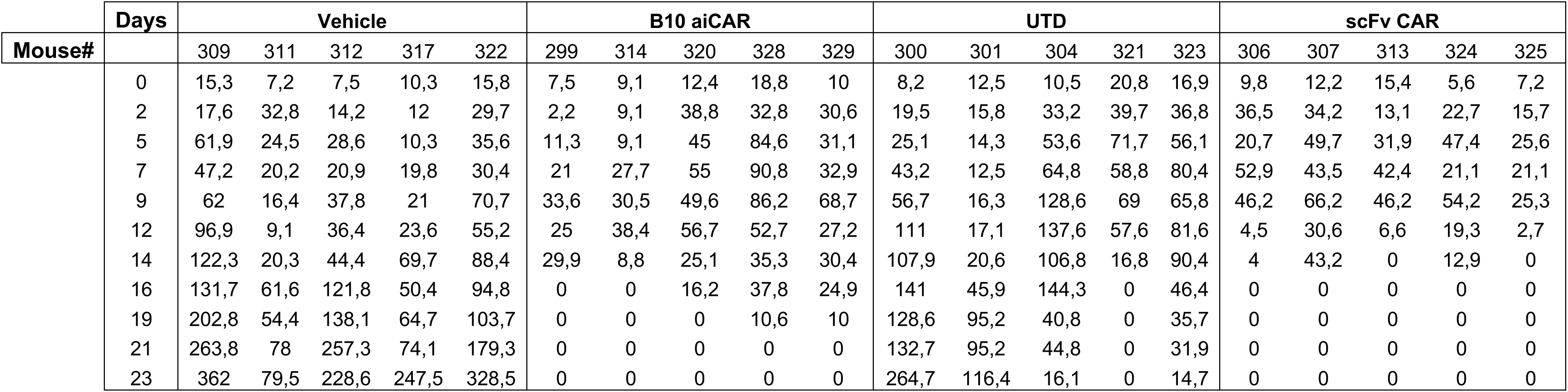
In vivo experiment tumor volumes (in mm. ^3^)

|  | Days | Vehicle |  |  |  |  | B10 aiCAR |  |  |  |  | UTD |  |  |  |  | scFv CAR |  |  |  |  |
| --- | --- | --- | --- | --- | --- | --- | --- | --- | --- | --- | --- | --- | --- | --- | --- | --- | --- | --- | --- | --- | --- |
| Mouse# |  | 309 | 311 | 312 | 317 | 322 | 299 | 314 | 320 | 328 | 329 | 300 | 301 | 304 | 321 | 323 | 306 | 307 | 313 | 324 | 325 |
|  | 0 | 15,3 | 7,2 | 7,5 | 10,3 | 15,8 | 7,5 | 9,1 | 12,4 | 18,8 | 10 | 8,2 | 12,5 | 10,5 | 20,8 | 16,9 | 9,8 | 12,2 | 15,4 | 5,6 | 7,2 |
|  | 2 | 17,6 | 32,8 | 14,2 | 12 | 29,7 | 2,2 | 9,1 | 38,8 | 32,8 | 30,6 | 19,5 | 15,8 | 33,2 | 39,7 | 36,8 | 36,5 | 34,2 | 13,1 | 22,7 | 15,7 |
|  | 5 | 61,9 | 24,5 | 28,6 | 10,3 | 35,6 | 11,3 | 9,1 | 45 | 84,6 | 31,1 | 25,1 | 14,3 | 53,6 | 71,7 | 56,1 | 20,7 | 49,7 | 31,9 | 47,4 | 25,6 |
|  | 7 | 47,2 | 20,2 | 20,9 | 19,8 | 30,4 | 21 | 27,7 | 55 | 90,8 | 32,9 | 43,2 | 12,5 | 64,8 | 58,8 | 80,4 | 52,9 | 43,5 | 42,4 | 21,1 | 21,1 |
|  | 9 | 62 | 16,4 | 37,8 | 21 | 70,7 | 33,6 | 30,5 | 49,6 | 86,2 | 68,7 | 56,7 | 16,3 | 128,6 | 69 | 65,8 | 46,2 | 66,2 | 46,2 | 54,2 | 25,3 |
|  | 12 | 96,9 | 9,1 | 36,4 | 23,6 | 55,2 | 25 | 38,4 | 56,7 | 52,7 | 27,2 | 111 | 17,1 | 137,6 | 57,6 | 81,6 | 4,5 | 30,6 | 6,6 | 19,3 | 2,7 |
|  | 14 | 122,3 | 20,3 | 44,4 | 69,7 | 88,4 | 29,9 | 8,8 | 25,1 | 35,3 | 30,4 | 107,9 | 20,6 | 106,8 | 16,8 | 90,4 | 4 | 43,2 | 0 | 12,9 | 0 |
|  | 16 | 131,7 | 61,6 | 121,8 | 50,4 | 94,8 | 0 | 0 | 16,2 | 37,8 | 24,9 | 141 | 45,9 | 144,3 | 0 | 46,4 | 0 | 0 | 0 | 0 | 0 |
|  | 19 | 202,8 | 54,4 | 138,1 | 64,7 | 103,7 | 0 | 0 | 0 | 10,6 | 10 | 128,6 | 95,2 | 40,8 | 0 | 35,7 | 0 | 0 | 0 | 0 | 0 |
|  | 21 | 263,8 | 78 | 257,3 | 74,1 | 179,3 | 0 | 0 | 0 | 0 | 0 | 132,7 | 95,2 | 44,8 | 0 | 31,9 | 0 | 0 | 0 | 0 | 0 |
|  | 23 | 362 | 79,5 | 228,6 | 247,5 | 328,5 | 0 | 0 | 0 | 0 | 0 | 264,7 | 116,4 | 16,1 | 0 | 14,7 | 0 | 0 | 0 | 0 | 0 |

## Methods

### Cell lines and mice

BxPC3, A549, K562, RPMI 8266, L-363, and Sup-T1 cell lines were obtained from the Leibniz Institute Deutsche Sammlung von Mikroorganismen und Zellkulturen (DSMZ), while HEK293T, SKOV3, and MM.1S cells were purchased from the American Type Culture Collection (ATCC). All cell lines were cultured under the conditions recommended by the provider company. For gene knock-outs in cell lines, the respective single guide RNA (sgRNA) (**Supplementary Table 2**; ordered as Alt-R CRISPR-Cas9 sgRNA, IDT) and Alt-R SpCas9 Nuclease V3 (cat. # 1081058, IDT) were pre-mixed at a molar concentration ratio of 3:1 (sgRNA:Cas9) to generate ribonucleoprotein (RNP) complexes. The target cell lines were electroporated with the RNPs using 4D-Nucleofector Core Unit & X Unit and the SF Cell Line 4D-Nucleofector X Kit S and the program DN-100. Subsequently, the cells were sorted for knock-out on BD FACSAria Fusion.

All animal studies were conducted under an Institutional Animal Care and Use Committee-approved protocol (Regierung von Oberbayern, Munich, Germany). Six-to eight-week-old female NOD.Cg-*Prkdc^scid^ Il2rg^tm1WjI^*/SzJ (NSG) mice were purchased from Charles River, Germany. NSG mice were subcutaneously injected with 5x10^6^ RPMI-8226 cells, and seven days later, after confirmation of engraftment, treated with a tail vain injection of vehicle, UTD, or 5x10^6^ CAR-T (all T cell treatments were normalized to the same number of total T cells). Tumor size was monitored by caliper measurement 3x/week, and tumor volume was calculated using the following equation: tumor volume [mm^3^]= (length × width^2^)/2. Mice were euthanized as specified in the animal study protocol, either at the end of the experiment or when meeting Institutional Animal Care and Use Committee predefined endpoints.

### TCE molecular cloning

All TCE constructs were cloned into a mammalian expression plasmid backbone under the control of a CMV promoter (following AgeI and NotI restriction digest of the parental plasmid; Addgene# 174828)^45^. The scFv sequences targeting EGFR and CD3 were obtained from publicly available sequences of cetuximab and OKT3 clone^46^. The EGFR-targeting ismb sequence was obtained from Cao et al., 2023^7^. The EGFR- and CD3-engaging domains were fused via a glycine-serine linker (GGGGS), and the entire TCE sequence was flanked by an Igκ leader peptide at the N-terminus and a His-tag element at the C-terminus. Gibson fragments with matching overlaps were PCR-amplified using Phusion high-fidelity polymerase (cat. # M0530L, NEB). The fragments were then gel-purified, Gibson assembled at 50 °C for 1 h, and transformed into chemically competent E. coli (cat. # C737303, Invitrogen). Gibson mix was made and used as previously described^47^. Plasmids used for transfection experiments were prepared using Qiagen Mini or Maxi Plus kits (cat. # 12123, # 12162, Qiagen).

### TCE production, purification, and quantification

HEK293T cells were seeded in a 6-well plate (7x10^5^) in DMEM containing 10% FCS. After 24h, the media of the cells was replaced with 2.5 ml Optimem (cat.# 31985062; ThermoFischer). The cells were transiently transduced with the indicated TCE constructs using a lipofectamine kit (cat. # L3000015; ThermoFischer) according to the manufacturer’s instructions. Briefly, 8ul lipofectamine in 2 ml Optimem was mixed with 1ug of plasmid DNA, 7ul P300 reagent in 2 ml Optimem, and incubated for 30 minutes at room temperature before transfecting HEK293T cells. After 24h, the supernatants containing the secreted TCE were collected. Supernatant from nontransduced HEK293T cells was used as a negative control.

### Flow cytometry-based measurement of TCE binding to target cells

To assess TCE binding to EGFR-expressing cells, TCE-containing supernatants were added (undiluted, or at the indicated dilutions) to BxPC3 cells and incubated for 2h at 4 °C. The bound TCE were then detected via flow cytometry using an anti-His-tag APC antibody (cat. # 362605, BioLegend).

### Flow cytometry, incl. list/table of all antibodies with catalog#

Cells were washed twice with FC-buffer (PBS + 2% FBS) and then incubated with antibody, diluted in FC-buffer, for 25 min at 4 °C in the dark. Next, cells were washed twice with FC-buffer and resuspended in FC-Buffer containing DAPI (cat. # 62247, Thermo Fisher) before being processed on BD FACSCanto II or BD LSRFortessa. Data were analyzed using FlowJo software version 10. The antibodies used can be found in **Supplementary Table 1**.

### Flow cytometry-based measurement of TCE binding to target cells

TCE were produced by transient transfection of the HEK293Tcells as described earlier. To assess their binding to EGFR-expressing cells, TCE-containing supernatants were added (undiluted, or at the indicated dilutions) to BxPC3 cells and incubated for 2h at 4 °C. The bound TCE were then detected via flow cytometry using a His-tag APC antibody (cat. # 362605, BioLegend).

### TCE-mediated cytotoxicity assay

Cryopreserved primary human T cells were thawed and cultured in RPMI medium (10%FCS and 1%P/S), and supplemented with 20 IU/ml recombinant human IL-2 (cat. #200-02; Peprotech) for 18h. T cells and BxPC3-CBG-GFP cells were resuspended at the indicated E: T ratios in Optimem supernatant containing TCE and co-incubated for 18h. Subsequently, a luciferase assay was performed as described above. The specific lysis of tumor cells mediated by TCE was calculated as follow: percentage of specific killing = [(target cells only relative luminescence units (RLU) − TCE RLU)/(target cells only RLU)] × 100.

### Design and cloning of CAR, target antigen, and luciferase constructs

All CAR, target antigen, and luciferase constructs were cloned into third-generation lentiviral vectors under the regulation of a human EF-1α promoter^19^. Vectors contain fluorescent reporter transgenes to assess transduction efficiency and cell viability in cytotoxic assays or to sort transduced cells. CARs carry a CD8 hinge, 4-1BB costimulatory domain, CD3 ζ signaling domain, and a fluorescent reporter (mCherry) following a self-cleaving peptide sequence (T2A). Constructs for target antigen expression feature the antigen of interest, a T2A-sequence, and a fluorescent reporter (GFP). For co-localization microscopy studies, some CAR and target antigen constructs were additionally cloned in a fused conformation; in these instances, the T2A sequence was replaced by a glycine-serine linker (GGGGS). A construct encoding click-beetle green luciferase and GFP separated by a T2A-sequence was generated for luciferase-based readouts. The nucleotide sequences were codon-optimized for expression in human cells or human T cells, respectively, using the GenSmart Codon Optimization online tool from GeneScript. Double-stranded DNA fragments of CAR binding domains, target, luciferase, and/or fluorescent reporters were ordered from IDT (Integrated DNA Technologies) and Twist Bioscience and assembled into the respective lentiviral plasmid backbone using Gibson assembly. Plasmids were then transformed into chemically competent E. coli (cat. # C737303, Invitrogen) and purified using Qiagen Mini or Maxi Plus kits (cat. # 12123, # 12162, Qiagen).

### CAR-T cell production

Primary human T cells were isolated from fresh human peripheral blood leukopaks (cat. # 70500.2, Stem Cell Technologies) or buffy coats (cat. # 350500, Bavarian Red Cross) obtained from healthy and anonymous donors using EasySep T Cell Isolation Kit (cat. # 17951, Stem Cell Technologies). For CAR-T cell generation, T cells were thawed at day 0, activated with anti-CD3/CD28 Dynabeads (cat. #11132D, Gibico) at a 1:3 ratio and cultured in RPMI 1640 medium supplemented with 10% fetal bovine serum (FBS), 1% penicillin-streptomycin and 20 IU/mL recombinant human interleukin-2 (IL-2) (cat. #200-02, Pepro Tech). On day 1, T cells were transduced with a CAR-encoding lentiviral vector at a multiplicity of infection (MOI) of 5 and expanded by medium doubling and IL-2 replacement every 2 days. After 5 days of expansion, Dynabeads were removed by magnetic separation. In parallel, activated but untraduced T cells (UTD) of corresponding donors were expanded and used for normalization and controls. On day 10, CAR-T transduction efficiency was determined by mCherry expression via flow cytometry analysis and normalized by adding UTD. For *in vitro* and *in vivo* experiments, CAR-Ts and UTDs were cryopreserved at day 10 and thawed for use in functional assays^19^.

### Luciferase-based killing for adherent cell lines and suspension cell lines

For luciferase-based killing assays, freshly thawed and normalized CAR-T or UTD were rested for 24h in RPMI 1640 with 10% FCS, 1% P/S supplemented with IL-2. Subsequently, the effector cells were co-cultured with luciferase-expressing tumor target cells at indicated effector-to-target (E:T) ratios at 37 °C and 5% CO2 for 16h. Cells were lysed with the Luciferase Cell Culture Lysis 5X Reagent (cat. # E1500, Promega) and further processed using the luciferase assay system (cat. # E1501, Promega) according to the manufacturer’s protocol. The luciferase activity was measured on a GloMax® Discover Microplate Reader (Promega) and provided as relative light unit values (RLU) by the GloMax® Discover System Software. Percentage specific lysis was calculated as follows: adherent cell lines: % specific lysis = (target cells alone RLU–total RLU with CAR T cells)/(target cells alone RLU) × 100%; non-adherent cell lines: % specific lysis = (total RLU with UTDs − total RLU with CAR-Ts)/(total RLU with UTDs) × 100%.

For adherent cell lines, tumor target cells were plated 24 h before adding the T cells. After 16h of co-culture, the T cells were removed by washing 4 times with PBS, and tumor cells were trypsinized, replated, and recovered for 2 days prior to cell lysis and luciferase activity measurement. For non-adherent cell lines, tumor cells and T cells were plated simultaneously, and tumor cells were lysed 16h later^19^.

### Proapoptotic protein and cytokine measurement (Legendplex assay)

Proapoptotic protein and cytokine release was measured in cell-free supernatants from 16h co-cultures of CAR-T and tumor cell lines at 1:1 E:T ratio and from 48h co-cultures of CAR-T and PDO at 10:1 E:T ratio using the Legendplex™ multi-plex bead-based immunoassay (cat. # 741187, Biolegend). For cytokine measurement, supernatants were diluted 1:1, and for proapoptotic protein measurement, supernatants were diluted 1:4 with ddH20, and the assay was carried out according to the manufacturer’s protocol.

### Cell avidity measurements

The protocol outlined here follows the most recent chip preparation guidelines provided by LUMICKS (v 17) as well as recently described methods^48^. The chips underwent activation with NaOH for 10 min at room temperature. Following two washes with ddH2O, the chips were then coated with Poly-L-Lysin (cat. # P4707, Sigma) for 10 min at room temperature. After coating, the chips were washed twice with ddH2O and once with warm PBS (cat. # 14190169, ThermoFisher). Then, they were flushed with serum-free RPMI1640 medium (cat. # 72400054, ThermoFisher) and incubated in a dry incubator at 37 °C until cell seeding.

SKOV3 and MM1.s cells were selected as target cells and matched with their corresponding chimeric antigen receptor (CAR) transduced Jurkat effector cells. All steps involving the target cells’ incubation were carried out in a dry incubator at 37 °C. SKOV3 target cells were seeded at a density of 100 x 10^6^ cells/mL, while MM1.s target cells were seeded at a density of 120 x 10^6^ cells/mL. After a 30-min incubation in serum-free RPMI1640 medium, a medium exchange was performed by substituting the medium with complete RPMI1640 medium, and the incubation was continued for 1.5 h. The chips were then returned to the dry incubator until measurement. CAR-expressing Jurkat effector cells were counted, washed once with warm PBS and stained (1 x 10^6^) with 1 µM cell trace far-red dye (cat. # C34572, Invitrogen) in 1 mL PBS for 15 min at 37 °C, 5% CO2. After incubation, the cells were washed once with warm complete RPMI1640 medium and adjusted to a concentration of 10 x 10^6^ cells/mL in complete RPMI for measurement. The stained effector cells were flushed into the z-Movi chips and co-incubated with the target cell monolayer for 15 min, until the application of gradually increasing force over 2 min and 30 s was started.

### Imaging the interaction of CAR-T and target cells

CAR-mCherry fusion constructs were used to transduce primary human T cells (MOI=5). After 10 days of expansion, live mCherry+ CAR T cells were sorted. K562 were transduced (MOI=1) with an EGFR-GFP fusion construct, and the live GFP+ cells were sorted ten days later. All sorted cells were cultured for one day to allow for recovery. Subsequently, CAR-T-mCherry cells (1x10^5^) and K562 EGFR-GFP cells (5x10^4^) were seeded onto a u-slide 8-well plate (cat. #80826; ibidi) in co-culture, in RPMI 1640 with 10% FCS, 1% P/S supplemented with IL-2 (20 IU/ml; cat. #200-02; Peprotech). To allow for settling and cell interaction, the cultures were incubated for one hour in an incubator (37°C, 5% CO2). Imaging was performed using a Leica Thunder Imager microscope containing an incubation chamber (37°C, 5% CO2). The images were captured at 63x magnification, with an optimal Z-stack step size. Additional image processing and cropping were performed with Leica Application Suite X (Las X version 3.8.1.26810), with the signal adjusted to avoid oversaturation. Image acquisition was performed in areas with sufficient CAR-T cell and target cell density (**Fig. 1e** and **Supplementary Data**).

### Long-term proliferation assay

The proliferative capacity of the CAR-T cells was assessed with repetitive antigen stimulation using EGFR- and BCMA-expressing and X-ray irradiated K562 cells as artificial antigen-presenting cells (APC). Normalized CAR-Ts and UTD were co-cultured with APC at a 1:1 E:T ratio, counted once weekly, and re-plated with APC.

### Generation and culture of PDAC patient-derived organoids

Primary patient-derived PDAC (pancreatic ductal adenocarcinoma) organoids were generated from either surgical resection or endoscopic ultrasound-guided fine needle biopsy (EUS-FNB). All reagents used for these experiments can be found in **Supplementary Table 3**. Surgical specimens were minced with a sterile scalpel and washed twice with Anti-Anti solution (Table 1). Biopsy samples were directly washed with Anti-Anti solution. Samples were then washed once with washing medium (Table 2) and digested in 5 mL of digestion medium (Table 3) using a falcon rotator at 37 °C and 30 rpm. To stop the enzymatic reaction, 15 mL of washing medium were added. The tissue pellets were incubated for 5 min with ACK lysing buffer/red blood cells lysis buffer (cat. # A1049201, ThermoFisher) and further digested with TrypLE (cat. # 12604039, ThermoFisher). The samples were washed with washing medium, and the cell pellets were resuspended in 200 µl of 10% BME (Cultrex Reduced Growth Factor Basement Membrane Extract, Type 2) (cat. # R3533-010-02, Pathclear) with PDAC PDO medium (Table 4) and 1:1000 Y-27632 (cat. # 10005583-10, Biomol/Cayman) in a 48-well plate. The use of patient material for this study was approved by the local ethics committee (Project 207/15), and written informed consent was obtained from the patients prior to the study.

### Live cell imaging of PDO CAR-T co-cultures

For live cell imaging, single PDAC cells obtained from PDO lines were seeded in 200 µl of 10% BME and PDAC PDO medium with 1:1000 Y-27632 in a 48-well plate. After 5 days of initial tumor cell seeding, CAR-T cells were added at a 10:1 ratio (effector cells: PDOs) on top of the wells in 200 µl of PDAC PDO medium with 1:1000 Y-27632. Live cell imaging was conducted using a Thunder Imager 3D Cell Culture (Leica Microsystems) immediately after co-culture, as well as at 12 h and 24 h post co-culture. Image acquisition was performed in areas with sufficient T cell and PDO density (**Fig. 1h**, **Extended Data Fig. 3a,** and Supplementary Data).

### RFdiffusion-based generation of BCMA binders

Generation of de novo protein binders was performed with the generative artificial intelligence (AI) tool RFdiffusion^9^ using the open-source Google Colab provided by Sergey Ovchinnikov. As the target protein, the B-cell maturation antigen (BCMA, PDB identifier: 1XU2)^30^ was chosen. RFdiffusion target contigs input was narrowed down by selecting a target chain in the PDB-identifier (1XU2, chain D). Optionally, target sites were then further specified by selecting suitable hotspots. The initial selection of hotspots was based on the known binding sites of the natural ligand APRIL. During the process of binder generation, some of these hotspots were then modified or supplemented with additional hotspots, to sample a more diverse binder set. The DNB length was set to 62 AA. For each BCMA binder design, at least 8, but up to 32 sequences were generated with ProteinMPNN, and the structure was afterward validated with AlphaFold2. Only AI-derived binders having triple helices as tertiary structures were selected for further evaluation. In the process of evaluating potential DNBs, we preferentially focused on the root mean-square deviation of atomic positions (r.m.s.d. value) based on the methods used to generate the previously published EGFR-targeting DNB^7^. Sequences with the lowest r.m.s.d. values were then selected for further evaluation and in vitro testing. Sequences and metadata of BCMA-targeting DNBs that were experimentally tested are listed in **Supplementary Table 4**. Visualizations of binder-target interactions were generated using PyMOL 3, and Fig. 1a was created using the Illustrate tool^49^.

### Teclistamab (TEC) and CAR-T cytotoxicity assays against BCMA-R27P

Freshly thawed primary human T cells or normalized CAR-Ts or UTDs were rested for 24h in IL-2 and subsequently used as effector cells. Primary human T cells were then co-cultured with K562 cell lines that express GFP as a fluorescent marker at a 5:1 E:T ratio with or without Teclistamab (Janssen) at varying concentrations (0.0001–100 nM)^36^. CAR-Ts or UTDs were co-cultured with the K562 cell lines at indicated E:T ratios. After 48h of incubation at 37 °C and 5% CO_2_ the co-cultures were prepared for flow cytometry analysis by staining for CD3 (cat. # 317318, Biolegend) and viability (DAPI; cat. # 62247, Thermo Fisher). Additionally, Precision Count Beads (cat. # 424902, Biolegend) were added following the manufacturer’s instructions to determine the absolute cell counts of living K562 cells. Using the BD LSRFortessa, 1x10^4^ Precision Count Bead events were collected per co-culture condition. After gating on living cells (DAPI negative) in the Indo-1 Violet channel and plotting the living cells in a two-parameter dot plot with APC in the vertical axis for CD3 expression T cells and FITC in the horizontal axis for GFP expressing K562 cells, the events of living K562 cells were determined by gating on APC-negative and GFP positive cells. The absolute amount of living K562 was calculated using the following formula: absolute cell count (cells/µl) = (event count K562 x Precision Count Beads volume) / (Precision Count Beads event count x final K562 flow volume). In order to compare K562 cell viabilities across different treatment conditions, the absolute K562 cell count was normalized to its respective control as follows: TEC cytotoxicity assay: % viability = (absolute cell count of K562 in the well of interest / absolute cell count of K562 only) x 100%. CAR-T cytotoxicity assay: % viability = (absolute cell count of K562 in co-culture of interest / absolute cell count of K562 in UTD co-culture) x 100%.

### Statistics and data reporting

Statistical calculations were performed using GraphPad Prism 10 and are described in the figure legends. Graphs were plotted using GraphPad Prism 10, protein structures were generated using PyMOL 3, and figures were assembled in Adobe Illustrator 28.4.

### Reporting Summary

Additional information on experimental designs, procedures, and analyses can be found in the Nature Research Reporting Summary.

## Data Availability

All raw images generated for the experiments shown in Fig. 1e, Fig. 1h, and Extended Data Fig. 3a can be found in the **Supplementary Data**. Plasmids encoding constructs used in this study will be deposited at Addgene.

## Code availability

RFdiffusion inputs and outputs (excluding trajectory files) that were used and generated during de novo binder design (see **Methods** and **Supplementary Table 4**) are provided in the **Supplementary Data**.

